# Characterization of sediment and granite hosted deep underground research laboratories reveals diverse microbiome functions, limited temporal variation and substantial genomic conservation

**DOI:** 10.1101/2024.03.28.587102

**Authors:** Yuki Amano, Rohan Sachdeva, Daniel Gittins, Karthik Anantharaman, Shufei Lei, Luis E. Valentin-Alvarado, Spencer Diamond, Hikari Beppu, Teruki Iwatsuki, Akihito Mochizuki, Kazuya Miyakawa, Eiichi Ishii, Hiroaki Murakami, Alexander L. Jaffe, Cindy Castelle, Adi Lavy, Yohey Suzuki, Jillian F. Banfield

## Abstract

Underground research laboratories (URLs) provide a window on the deep biosphere and enable investigation of potential microbial impacts on nuclear waste, CO_2_ and H_2_ stored in the subsurface. We carried out the first multi-year study of groundwater microbiomes sampled from defined intervals between 140 and 400 m below the surface of the Horonobe and Mizunami URLs, Japan. We reconstructed draft genomes for >90% of all organisms detected over a four year period. The Horonobe and Mizunami microbiomes are dissimilar, likely because the Mizunami URL is hosted in granitic rock and the Horonobe URL in sedimentary rock. Despite this, hydrogen metabolism, rubisco-based CO_2_ fixation, reduction of nitrogen compounds and sulfate reduction are well represented functions in microbiomes from both URLs, although methane metabolism is more prevalent at the organic- and CO_2_-rich Horonobe URL. High fluid flow zones and proximity to subsurface tunnels select for candidate phyla radiation bacteria in the Mizunami URL. We detected near-identical genotypes for approximately one third of all genomically defined organisms at multiple depths within the Horonobe URL. This cannot be explained by inactivity, as in situ growth was detected for some bacteria, albeit at slow rates. Given the current low hydraulic conductivity and groundwater compositional heterogeneity, ongoing inter-site strain dispersal seems unlikely. Alternatively, the Horonobe URL microbiome homogeneity may be explained by higher groundwater mobility during the last glacial period. Genotypically-defined species closely related to those detected in the URLs were identified in three other subsurface environments in the USA. Thus, dispersal rates between widely separated underground sites may be fast enough relative to mutation rates to have precluded substantial divergence in species composition. Species overlaps between subsurface locations on different continents constrain expectations regarding the scale of global subsurface biodiversity. Overall, microbiome and geochemical stability over the study period has important implications for underground storage applications.

## Introduction

The subsurface, especially hundreds of meters below ground, is one of the last biological frontiers (Teske et al., 2013; Hoshino et al., 2020). Beyond basic scientific interest in organisms that live there, the microbiology of these regions is an important practical consideration if they are to be used to store nuclear waste and other materials (e.g., CO_2_ or H_2_). Microbes in proximity to underground repositories may impact the stability of the stored material via container corrosion (e.g., Pedersen 1999; Rajala et al., 2015; Stroes-Gascoyne et al., 2010; Stroes-Gascoyne and West, 1997), by consumption of stored resources (e.g., H_2_, Dopffel et al., 2023; Liu et al., 2023) or by degrading containment integrity (e.g., via increasing rock porosity and/or permeability, Zeng et al., 2023).

Much of what we know about subsurface life (below the soil zone) comes from studies of groundwater. Genome-resolved microbiome studies of 3 - 6 m depth groundwater in an aquifer adjacent to the Colorado River, Rifle, Colorado brought to light dozens of new lineages of bacteria and archaea (Wrighton et al., 2012; Anantharaman et al., 2016). Organisms from these groups have since been described from numerous other ecosystems, including northern California groundwater aquifers (He et al. 2021) and groundwater delivered to the surface via eruption of a cold CO_2_-driven geyser (e.g., Probst et al., 2018). However, these systems provide limited insights into deep subsurface microbiology. Recovery of deep underground samples for biological analysis is possible via drilling, but drilling provides samples with little context, interpretation of the results may be complicated by contamination (e.g., from drilling fluids), and samples represent only a single time point (Mouser et al. 2016). For this reason, the construction of large, human-accessible deep subsurface research laboratories has been an important development. Within these underground research laboratories microbiological samples can be collected from well defined sites with minimal contamination. Important examples of such laboratories include Äspö in Sweden (Hallbeck & Pedersen, 2008; Mehrshad et al., 2021), Grimsel Test Site in Switzerland (Konno et al., 2013), the Mont Terri Underground Rock Laboratory in Switzerland, the Horonobe Underground Research Laboratory (Horonobe URL) in Hokkaido, Japan and the Mizunami Underground Research Laboratory (Mizunami URL) in Gifu, Japan. Here, we conducted the first multi-year investigation of subsurface microbial diversity and metabolic capacities in microbiomes of the two Japanese URLs. This work follows one prior publication that considered microbial metabolisms in some Horonobe URL microbial communities (Hernsdorf et al., 2017) and another that ecologically and genomically profiled anaerobic methane-oxidizing archaea in the Mizunami URL (Ino et al., 2017). Our analyses provide a comprehensive genomics-based overview of microbial and metabolic diversity and spatial and temporal variation. We uncover genotypic overlap within each URL and likely geologically-based differences between the microbiomes of the URLs. Finally, we compare genotypes from the URLs to genotypes of bacteria and archaea sampled from three underground locations in the USA. The results begin to address the ‘(to what extent is) everything everywhere’ question for microbiomes that exist way below the Earths’ surface, thus providing clues to overall levels of subsurface microbial diversity.

## Results

Our research was conducted primarily in two underground research laboratories (URLs) in Japan (**Figure 1a**). From the Mizunami URL (35°22’40.68”N, 137°14’15.63”E) we analyzed metagenomic data from 7 groundwater samples that were recovered from 200 to 400 m below the surface (**Table S1**; **Figure 1b**). During URL construction, 4-7 years prior to the first sample collection in 2014, ∼10 cm diameter, ∼100 m long boreholes were established by coring into rock from tunnel walls. At six locations in three access tunnels (200 m, 300 m, 400 m below the surface), water was extracted from sampling zones at 26.9 - 96.1 m distances from the tunnels (**Table S2**). The seventh sample was collected from the 200 m depth site in 2015. Low ionic strength groundwater in these regions is organic-poor, Na^+^-Ca^2+^ - Cl^−^ brine (Iwatsuki et al., 2005; Hayashida et al., 2016), with temperatures ranging between 15 and 23 °C. The pH ranges from 8.7 to 9.1 and Eh values range from −180 to −60.7 mV (**Table S3**). From the Horonobe URL (45°02’41.92”N, 141°51’34.20”E) we analyzed metagenomic data from 19 groundwater samples collected from five locations at 4 - 57 m from the tunnels (**Table S1; S2; Figure 1c; Figure S1**). Groundwater was recovered from 140 m and 250 m below the surface via tubes into defined intervals of boreholes into the surrounding rock. The Horonobe groundwater is organic rich and saline, dominated by Na^+^-Cl^−^ - HCO ^−^ (**Table S3**, Amano et al., 2012; Sasamoto et al., 2015). Temperatures range from 13 - 25 °C, pH ranges from 6.4 to 7.2, and Eh values range from −315 to −204 mV. Unlike at Mizunami, the Horonobe groundwater is saturated with both CH_4_ and CO_2_ (Miyakawa et al., 2017; Tamamura et al., 2018), which degas during sample recovery.

**Figure 1.**
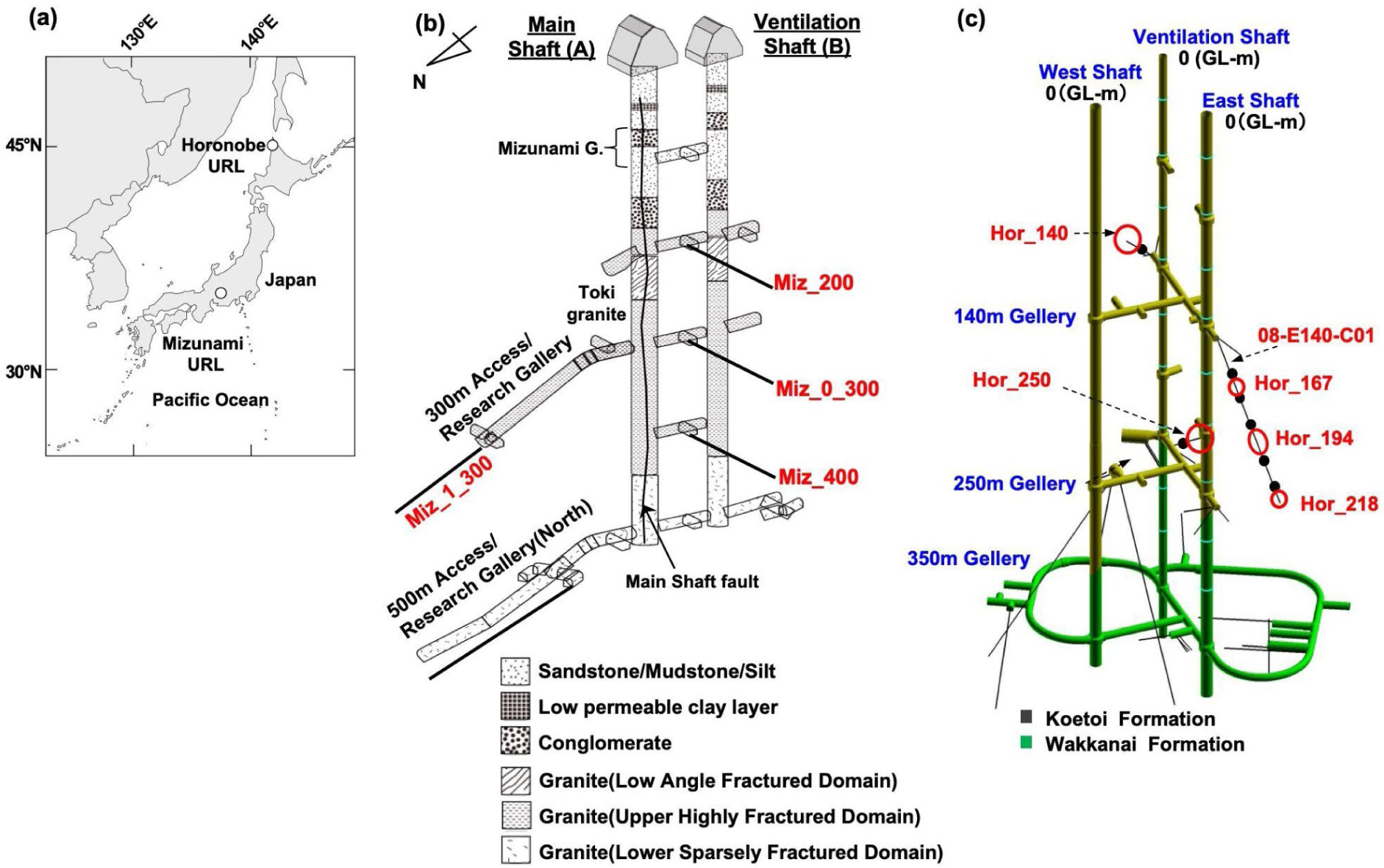
(**a**) Map of the Mizunami and Horonobe URL locations in Japan. (**b**) Layout of boreholes in shafts and galleries in the Mizunami URL and (**c**) the Horonobe URL.

Total DNA was extracted from the 26 samples and between 10.6 and 16.4 Gbp of 150 bp paired end Illumina sequences obtained from each sample. The sequences from each sample were assembled independently and genomes reconstructed. After gene prediction and functional annotation, a census of organisms was performed based on phylogenetic classification of ribosomal protein S3 (rpS3) sequences. The abundance of each organism was determined based on read coverage values for the rpS3-bearing scaffolds. We reconstructed 1154 rpS3 sequences from the Horonobe metagenomes and 624 from Mizunami metagenomes. Phylogenetic analysis revealed that the vast majority of rpS3 sequences from both the Mizunami and Horonobe URLs were from Candidate Phyla, major groups that lack even a single isolated representative (**Figure 2; Data S1**). Up to 21 % of Mizunami rpS3 sequences place within the DPANN. No DPANN sequences were recovered from the Horonobe URL, although Altiarchaeota are present (up to 90% of sequences) and they are sometimes grouped with DPANN (Adam et al., 2017). Up to 47% of Mizunami sequences and up to 41% of Horonobe sequences are from bacteria of the Candidate Phyla Radiation (CPR; Brown et al. 2015; **Figure 3, 4**).

**Figure 2.**
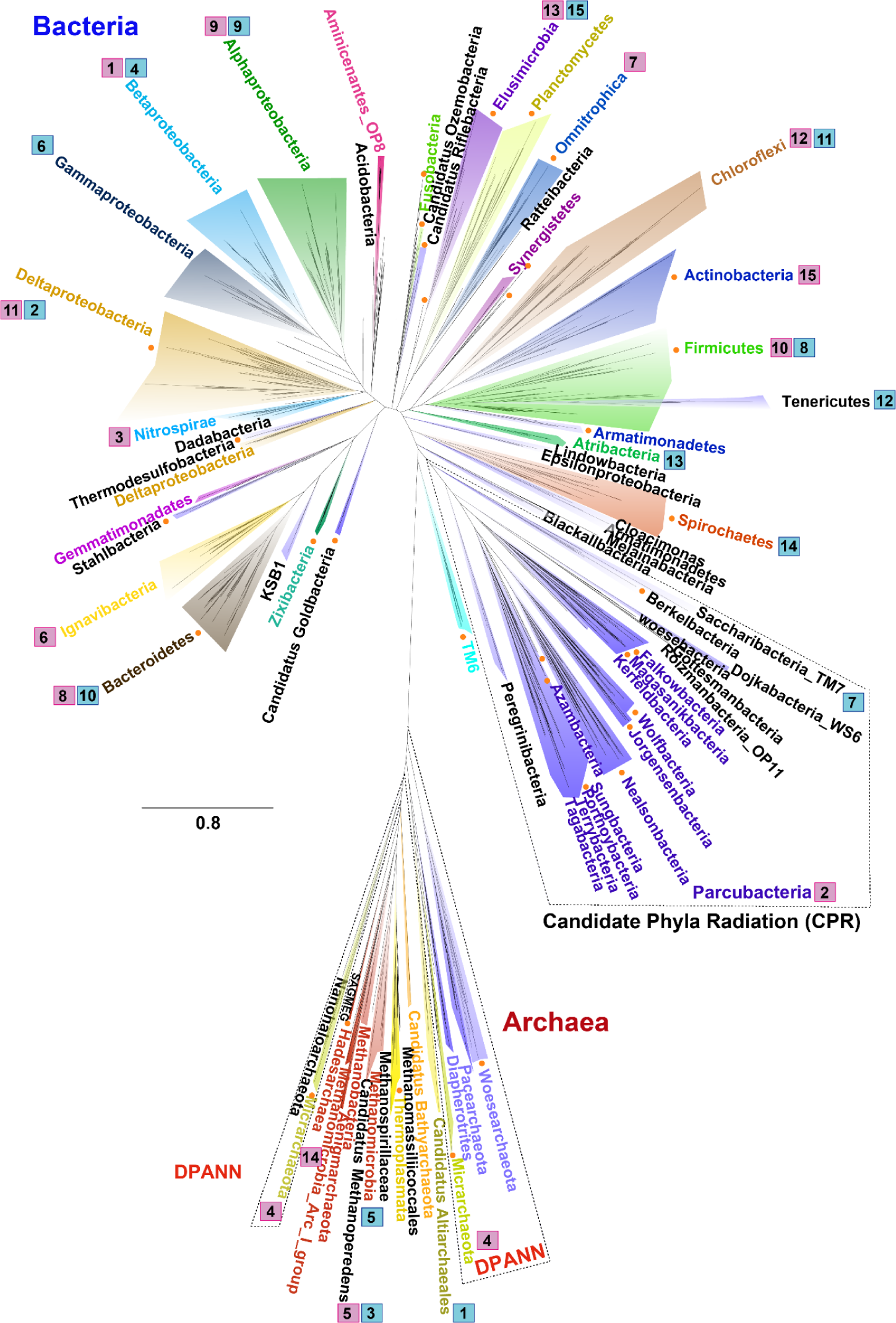
Phylogenetic tree of 956 representative sequences of ribosomal protein S3 from the Horonobe and Mizunami samples, along with reference sequences. Orange dots indicate individual sequences with <70% rpS3 amino acid identity to sequences in NCBI. Numbers in squares show the average rank abundance of each phylum (or class in the case of Proteobacteria). Blue squares indicate organisms from the Horonobe URL and pink squares indicate samples from the Mizunami URL.

**Figure 3.**
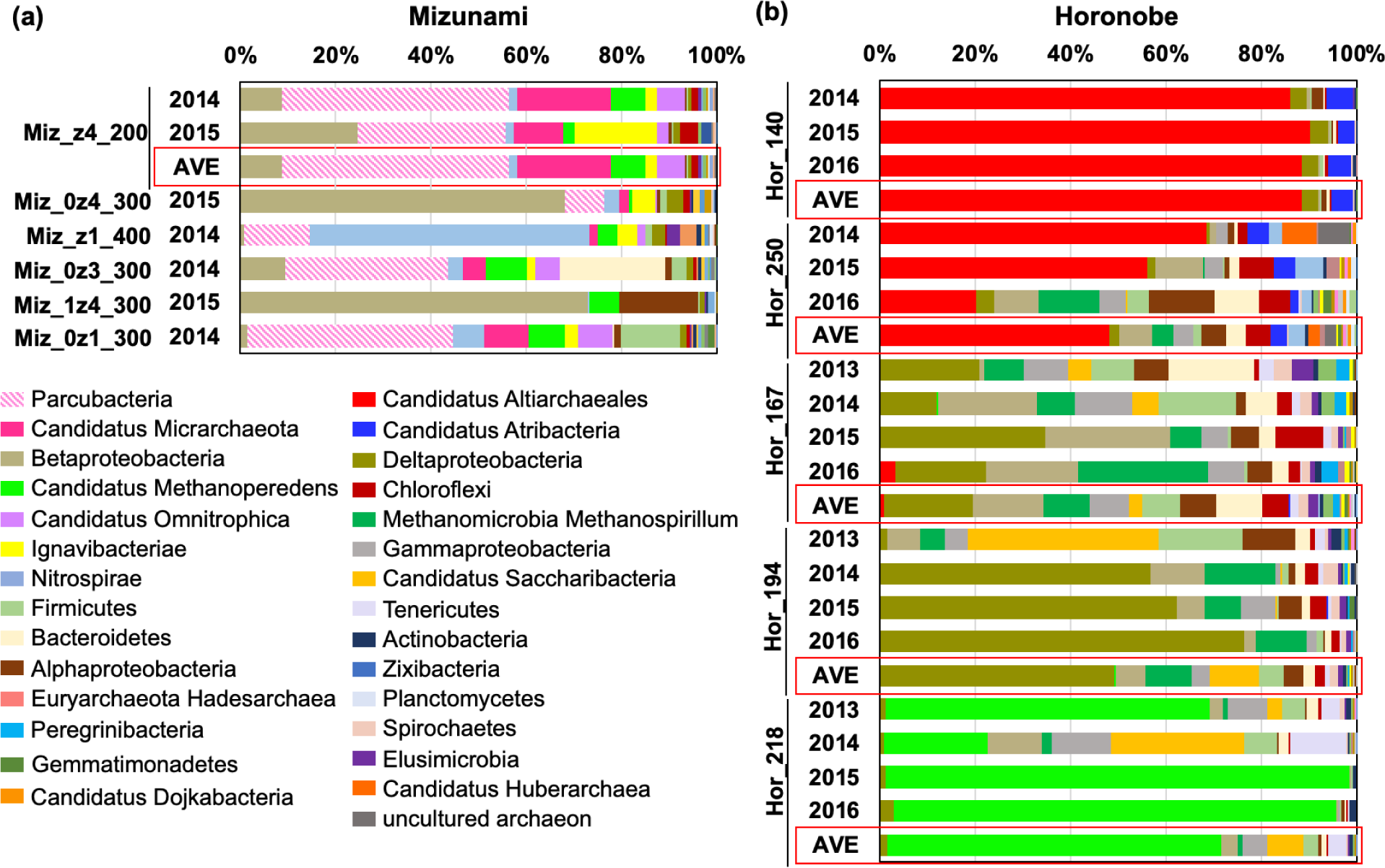
Overview of URL microbial diversity based on the 15 most abundant organisms in each sample, classified mostly at the phylum/class level. Samples are listed in order of increasing the distance between the sampling site and the closest access tunnel to seek evidence of perturbation due to the presence of the tunnel. (**a**) 7 Mizunami 0.2 µm-filter URL samples collected between 2014 and 2015. (**b**) 18 Horonobe URL 0.2 µm-filter samples collected between 2013 and 2016. Given the general consistency in community composition over time, we also averaged the compositional data from the same site for subsequent analyses (red boxes).

**Figure 4.**
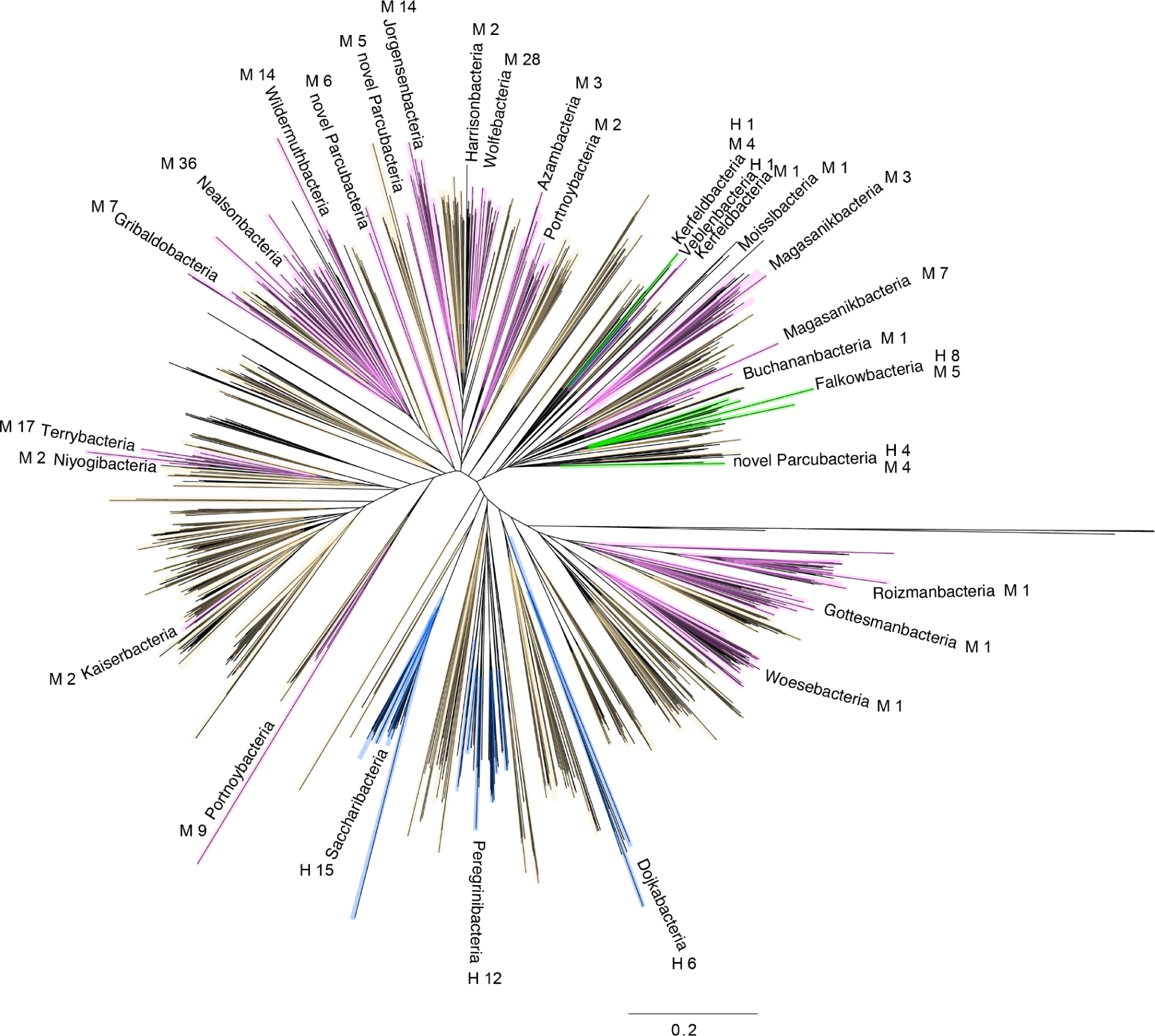
Phylogenetic tree for CPR bacteria constructed using ribosomal protein S3 sequences. Pink and blue shadings indicate sequences from the Mizunami and Horonobe URLs, respectively. Green branches indicate lineages with sequences from both the Mizunami and Horonobe URLs. The numbers after M (Mizunami) and H (Horonobe) indicate the number of sequences in each named lineage. Brown highlighted sequences are reference sequences. The long branches without color indicate Archaea, which were used as the outgroup.

Initially, we profiled community composition over space and time using high level (mostly phylum) phylogenetic groupings (**Figure 3, Figure S2**). However, for the Horonobe URL we also evaluated class level diversity and determined that the phyla with many species representatives also have species assigned to more classes, predicted by a linear trend (slope of 0.39; **Figure S3a)**.

We hypothesized that URL construction may have perturbed, and may continue to perturb, the microbiomes and thus community composition may correlate with distance from the access tunnels. Despite relatively consistent geological settings within each URL there are substantial differences in phylum-level community composition between some sites. Within the Mizunami URL, the abundances of Parcubacteria and Micrarchaeia (DPANN) vary substantially, and they were essentially absent from one 300 m depth site in a low-fluid flow zone. In the Horonobe microbiomes, Altiarchaeales (SM1) were abundant only in the two sites (140 m and 250 m) located very close to the URL tunnels. Horonobe samples collected at intermediate distances from the tunnels featured abundant *Deltaproteobacteria*, *Methanomicrobia, Firmicutes, Betaproteobacteria*, and *Bacteroidetes*. Candidatus Saccharibacteria was abundantly detected only in 2013.

*Methanoperedens* (ANME-2d) were prevalent at the site most distant from the tunnels. Parcubacteria and other CPR bacteria were abundant in samples collected from high fluid flow zones at distance from the tunnels. Despite site to site differences, Horonobe samples collected over three or four years from the same sites showed little variation in microbiome composition (**Figure 3b**).

Given the difference in surrounding rock type between the two URLs, we were interested to know how similar the microbiomes of the granitic hosted Mizunami URL are to those of the sediment-hosted Horonobe URL. The similar microbiome compositions within each sampling location over time made it reasonable to average phylum compositions at each location to enable comparison of the URLs at a high taxonomic level (**Figure 3**). For both URLs, the 15 most abundant organisms are from 15 different major (mostly phylum-level) groups and the URLs share only 8 of these (Firmicutes, Alphaproteobacteria, Bacteroidetes, Chloroflexi, ANME-2d (Methanoperedenceae), Betaproteobacteria and Elusimicrobia). Deltaproteobacteria were far more abundant in Horonobe compared to Mizunami samples. Nanoarchaeal Micrarchaeota are only prevalent in the Mizunami URL whereas Altiarchaeales archaea (SM1) and Saccharibacteria (TM7) are only prominent in the Horonobe URL (**Figure 2,3; Table S4)**. The CPR bacterial types present in Horonobe samples (Peregrinibacteria, Saccharibacteria and Dojkabacteria) are different from those in the Mizunami samples (primarily diverse Parcubacteria groups) (**Figure 4; Table S4**). Given these results and the relative consistency of groundwater microbiology over time, we conclude that the microbiomes of the Mizunami and Horonobe URLs are distinct at high taxonomic levels. We also evaluated microbial overlap between the URLs at the species level. Only 15 out of 490 species occur in both URLs (3 Actinobacteria, 1 Bacteroidetes, 1 Betaproteobacteria, 4 Deltaproteobacteria, 1 Firmicutes, 2 Elusimicrobia, 1 Ignavibacteria, 1 Spirochaete and 1 Candidatus Kuenenbacteria).

Dereplication of the 225 draft genomes from the Mizunami URL and 265 draft genomes from the Horonobe URL yielded 489 genomes in total, implying just one genomically defined organism (Clostridia, Species 4) was shared between the two URLs. The reconstructed genomes account for 93% and 90% of all organisms detected at the Mizunami and Horonobe URLs, respectively, based on rpS3 analysis. Organisms lacking a draft genome occur in only a small subset of samples (for Mizunami, all were detected in only one sample). Thus, for both URLs, we conclude that the microbiomes of both URLs are now extensively genomically sampled.

Analysis of the presence/absence patterns of all species across samples (**Figure 5**, **Table S5)** showed that many organisms are present in multiple sites within each URLs. One organism, a *Rhodobacterales,* occurred in 16 of the 19 Horonobe sampling sites. Hierarchical clustering of the presence absence patterns for the Horonobe URL (**Figure S3b)** showed that community compositional similarities are well predicted by the sampling site (i.e., samples collected at the same site in different years cluster together). Clustering of Horonobe samples based on organism presence/ absence patterns supports the hypothesis that microbiome distance of the access tunnels may impact the organisms present.

**Figure 5.**
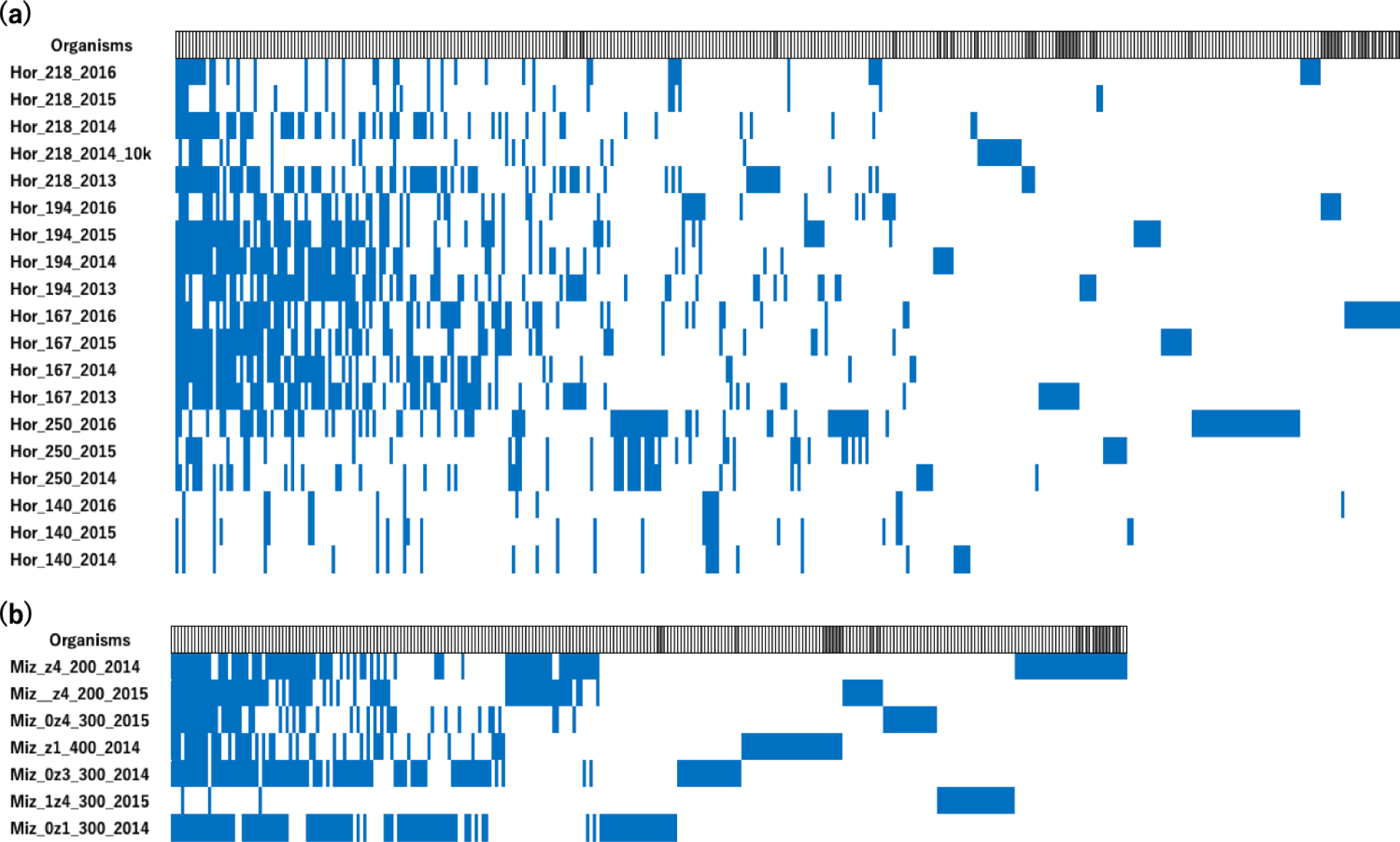
Detection (blue bars) of organisms (columns) in samples (rows) listed in approximate order of decreasing distance from the access tunnels. Organisms lacking genomes are indicated by a dark gray box in the Organisms bar. (**a**) Almost one quarter of all organisms detected within the Horonobe URL were present in >25% of the samples. 90.389.2% of all organisms are represented by draft genomes. Organisms lacking genomes were all detected in ≤ 3 samples. (**b**) Within the Mizunami URL, 45% of organisms were detected in at least 25% of the samples and 10% of all organisms detected were present in >70% of the samples. 92.0% of organisms are represented by draft genomes. All organisms lacking genomes were detected in just one sample. For details see **Table S3**.

The extensive genomic sampling of both URL microbiomes made it reasonable to use the genomes to provide an overview of biogeochemical capacities in these subsurface ecosystems. Our analysis targeted genes involved in CO_2_ fixation, hydrogen cycling, nitrogen cycling and sulfur cycling. For example, we analyzed the diversity of Rubisco large subunit sequences, focusing primarily on forms I and II that co-occur with phosphoribulokinase, as these genes are indicative of the capacity for CO_2_ fixation via the CBB cycle (**Figure S4; Table S6**). Genes for this pathway were abundant, and detected in all samples of both URLs. Hydrogenase-encoding genes were recovered from all the samples from both URLs, indicating a broad distribution (**Table S7**). Most of the hydrogenases in Mizunami were assigned to groups 1a, 3b, 3c, 3d, 4g [NiFe]-hydrogenases and groups A3 and C3 [FeFe]-hydrogenases. Interestingly, the most abundant [NiFe]-hydrogenase enzymes in Mizunami URL are from group 3b, which is predicted to oxidize NADPH and evolve hydrogen, maybe reversibly (Greening et al., 2016). These complexes may have sulfhydrogenase activity (elemental sulfur (S^0^) reduced to polysulfide to H_2_S; Ma et al., 1993). Group 1a [NiFe]-hydrogenase is predicted to oxidize H_2_ and is predominantly found in anaerobic Firmicutes and Deltaproteobacteria capable of H_2_-dependent sulfate reduction and metal reduction. On the other hand, the hydrogenases from the Horonobe URL were from all of the four major [NiFe] groups and the three major [FeFe] groups. The abundant [NiFe]-hydrogenases were assigned to groups 3b, 3c, 3d, and 4f. Group 4 [NiFe]-hydrogenases are predicted to be membrane-bound H_2_-evolving enzymes. Although the function of group 4f is uncertain, it may couple oxidation of a one-carbon compound to proton reduction concurrent with proton translocation. Overall, the indications are that H_2_ is an important energy currency in these anoxic subsurface environments.

Abundant genes for nitrogen cycling were identified in genomes from both URLs (**Table S8**). These include genes involved in nitrogen fixation, nitrate reduction, nitrite reduction to ammonia and nitric oxide reduction. In the Mizunami URL, up to 53% of genomes encode genes for nitrite reduction to ammonia compared to up to 13% in the Horonobe URL. No genes for oxidation of ammonia were detected in any sample from either URL, which may explain the very high concentrations of ammonia in the groundwater at Horonobe (Sasamoto et al., 2018). At Mizunami, a widely distributed *Rhodocyclales* has the capacity for nitrate reduction, nitrite reduction to ammonia and sulfate reduction. The *Rhodobacterales* that is the most widely distributed organism within the Horonobe URL also has genes for reduction of nitrate, nitrite reduction, nitric oxide and sulfur cycling. Similar capacities are predicted for other widely distributed bacteria (e.g,. Gammaproteobacteria).

Genes for sulfate reduction, sulfite reduction, sulfur oxidation, and thiosulfate oxidation, were observed in genomes of organisms from both URLs, but are more prevalent in the genomes of organisms from Mizunami compared to Horonobe (**Table S8**). Abundant genes for sulfur dioxygenase (sdo), which could be involved in oxidation of elemental sulfur to sulfite, were detected in both URLs. Genes for sulfate reduction were particularly prevalent in most *Nitrospirae* and some Deltaproteobacteria from the Mizunami URL and diverse Deltaproteobacteria from the Horonobe URL. However, the two most abundant *Nitrospirae* associated with the Mizunami URL apparently lack this capacity. Interestingly, genes for selenate reduction were prevalent in genomes of Deltaproteobacteria and Spirochaetes from both URL microbiomes. In fact, selenate reduction is the only biogeochemically relevant capacity predicted for a Spirochaete that occurs in 15 of the 19 Horonobe sampling sites.

We asked how similar the collections of genomically defined organisms from Mizunami and Horonobe were to groups of organisms from other genomically well sampled subsurface groundwater systems. Our comparison set comprised organisms from the two URLs, from the cold, CO_2_ driven Crystal Geyser that taps into deep aquifers below the Colorado Plateau (Probst et al. 2018), the shallow groundwater aquifer adjacent to the Colorado River, Rifle, Colorado (Anantharaman et al. 2016), and groundwater from northern California (He et al. 2021). We chose these due to their high genomic resolution and, in part, because the metagenomic datasets were processed using essentially the same protocols as the current study. We did not include comparisons to soils, as there have been recent reviews that show that soils worldwide are dominated by Acidobacteria, Actinobacteria and Gemmatimonadetes, groups generally not prominent in the URL environments. We used each of the 1472 rpS3 sequences from genomes from the five locations to approximate species and compared levels of species overlap between locations. Notably ∼45% of the Horonobe sequences are most similar to sequences from the sediment-hosted Rifle aquifer **(Table S9**).

To minimize the effect of substantial differences in the number of genomes reconstructed from each location on comparisons, we also evaluated species-level similarity based on average % amino acid identity (aa% ID) of the best matching sequences for each rpS3 protein (**Figure 6**). Notably, the northern California groundwater genome set shows statistically significant differences in similarity in all comparisons (i.e., more similar to the set from Rifle than Mizunami, more similar to the set from Mizunami than from Horonobe, etc.). The genomes from Horonobe are more similar to those from Rifle compared to northern California groundwater (p ≤ 0.05) and less similar to Mizunami, although the statistical significance is just above our cutoff (p > 0.05).

**Figure 6:**
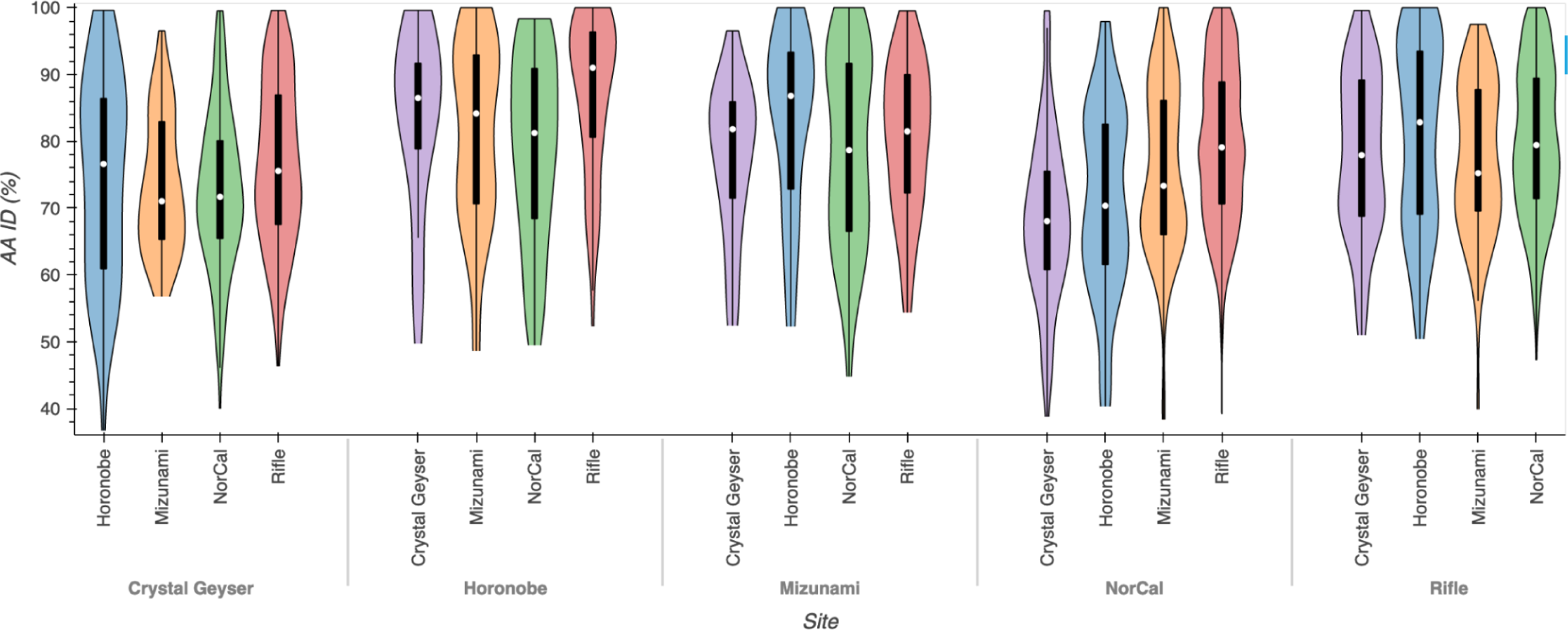
Comparison using the dereplicated genome sets from five genomically well sampled terrestrial subsurface ecosystems to seek instances where two ecosystems are similar or significantly different from each other. The highest scoring pairwise hit for each of 1472 sequences from genomes was assigned to an ecosystem comparison category and the aa ID (%) inventoried. For example, when all 503 sequences from Crystal Geyser were compared to all sequences from the four comparison datasets, there were 59 instances where the closest sequence was found in the Horonobe dataset. The 59 aa ID % values are represented by column 1. Of the comparisons, those that were significantly different are Horonobe compared to northern California (NorCal) vs. Rifle (p value of 1 x 10^−3^), Crystal Geyser compared to Rifle vs. NorCal (p value of 5 x 10^−3^), and NorCal compared to all other ecosystems (p < 1 x 10^−5^. For details see **Table S5**.

We evaluated genome novelty at all five locations by considering only those organisms whose rpS3 protein shared < 50% aa% ID with the highest scoring sequence from a comparison location. The most divergent genomes sampled from the Horonobe URL are for Hadesarchaeota, Thermoplasmatales or Bathyarchaeota, and from the Mizunami URL, were DPANN archaea. Notably, ∼80% of most divergent sequences across the five datasets were from Archaea (almost half of these are DPANN) and almost half of the bacterial cases were CPR bacteria.

Generally lacking to date have been analyses that leverage genome collections to investigate the extent of genomic overlap of highly related (i.e., rpS3 proteins with > 99% aa% ID) organisms across regions distant from each other geographically (e.g., different continents) and/or physically (e.g., due to depth below the surface). We sought cases where highly related organisms were shared between subsurface locations in the USA and the two Japanese URLs and found 106 shared bacteria and archaea from 16 different classes/phyla (Table S10). Most of the pairs were from Horonobe and Rifle or northern California groundwater and Rifle. Eight genomes with identical rpS3 protein sequences (from 6 different phyla/classes) are classified as representing the same species based on >95% genome average nucleotide identity (**Table S10**). One Bacteroidetes species (genus *Lentimicrobium*) from Horonobe (detected from 167 m, 218 m and 250 m samples) shares 99 % ANI with a genome from the Rifle aquifer. The *Lentimicrobium* genomes are nearly perfectly syntenous across aligned regions **(Figure S5**), consistent with these organisms being very closely related. Thus, we conclude that there is some strain-level overlap between geographically distant shallow and deep subsurface regions.

To put the results of this multi-location comparison into context we turned back to the Horonobe dataset, which has sufficient sampling to provide insight into genomic similarity for organisms sampled from the same site at different times and from different depths and locations. We identified 99 distinct genome clusters (451 genomes), each of which is comprised of genomes that share > 99.5% ANI over > 95% of the genome alignment. Of the 99 Horonobe clusters, 17 consisted only of organisms from the same site sampled in different years. For example, a Peregrinibacterium for which we manually curated a 998,424 bp complete genome (167 m sample) shares 99.997% ANI over the entire genome with a genome collected a year later (and 100% ANI over 993 kbp) from the same site (this genotype was detected in all four years). The remaining 83% of clusters, containing members from different depths within the URL, represent 31% of all genomes reconstructed. Organisms from the shallowest depth (140 m) were least commonly clustered with organisms from other depths. However, when they were clustered, it was typically with organisms from the deepest site (250 m). Approximately one third of the clusters contain organisms from three different depths and 12% contain organisms from the four deeper sites. Strikingly, some clusters included genomes from different sites with average pairwise ANI values of >99.997 %. For example, clusters of genomes for *Saccharibacterium*, *Clostridium, Syntrophobacterales* and a *Gammaproteobacteria* from four different depths all have ≥ 99.98% pairwise ANI. The complete *Peregrinibacteria* genome (167 m sample) is identical to a genome comprised of three contigs from the 194 m depth site (98.8% genome alignment). Thus, we conclude that extremely closely related bacteria occur at sites separated by large volumes of rock.

We considered the possibility that the very closely related species from different depths at the Horonoble URL may be a distinct, possibly inactive subset of the microbial communities. However, the taxonomic affiliations of the widely distributed species is well predicted by the taxonomic composition of the overall Horonobe genome set. Thus, we considered the possibility that the genomic homogeneity of species throughout the Horonobe URL is due to overall microbiome inactivity. We tested for in situ growth by calculating an index of replication (iRep; Brown et al. 2016) for a subset of organisms present at a variety of abundance levels. The Peregrinibacteria genome that was highly conserved in the 167 m and 194 m samples had an iRep value of 1.14, similar to values for other abundant microbes affiliated with the Firmicutes (1.24), Methylophilales (1.12) and Rhodobacterales (1.26). Low coverage values limited the set of genomes for which this calculation was possible.

**Figure 7.**
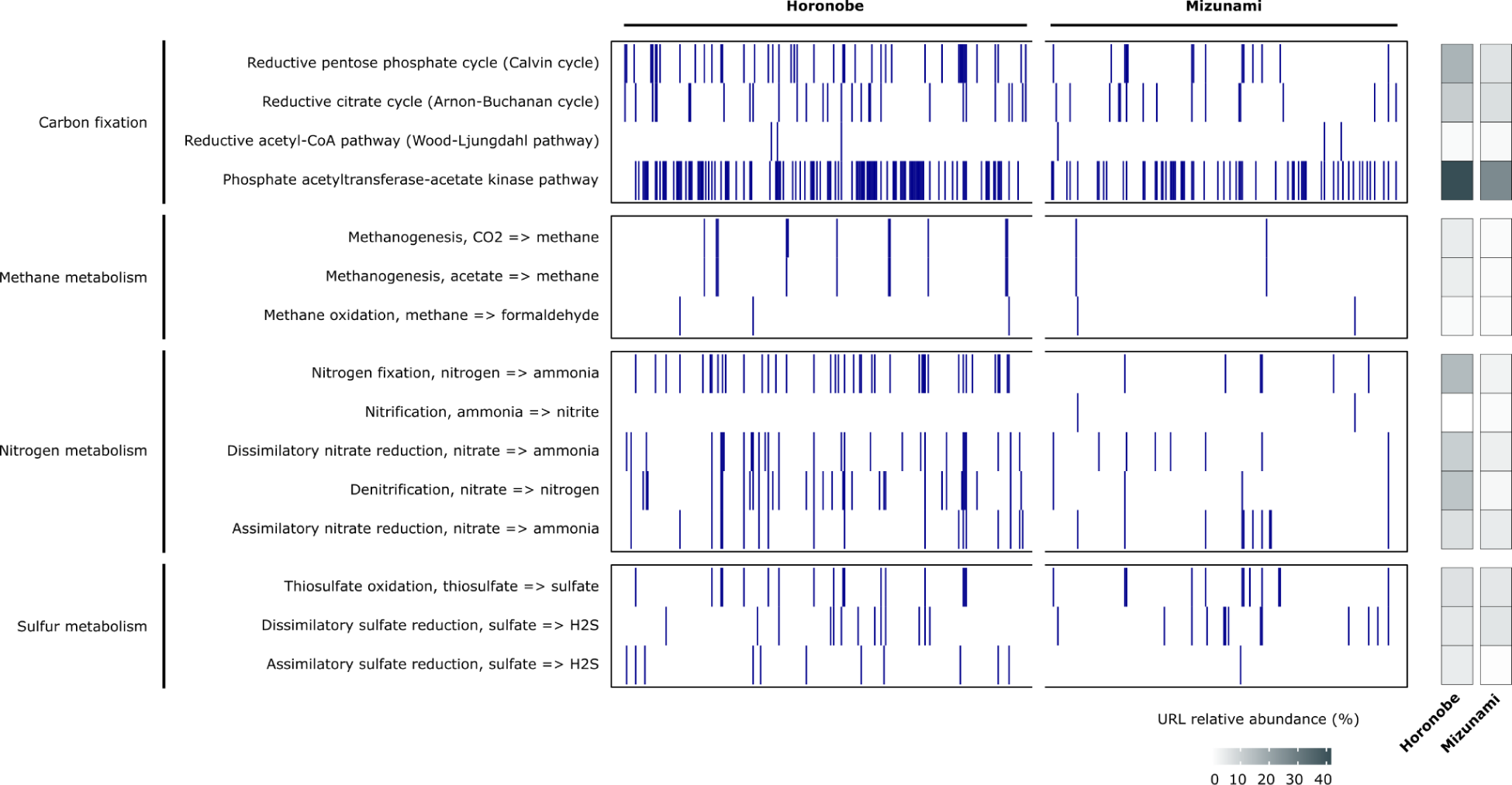
Key metabolisms across the Mizunami and Horonobe URLs. Presence/absence of each metabolic pathway based on the occurrence of indicative marker genes annotated with KEGG Orthology using Kofamscan (y-axis) in each of the recovered genomes (x-axis). The URL relative abundance (%) shows the proportion of the 265 Horonobe and 225 Mizunami genomes with that metabolism.

## Discussion

Underground research laboratories provide unique access to the deep subsurface biosphere. Findings from such laboratories are relevant for understanding Earth’s microbiomes broadly, and may inform plans to use the subsurface for storage of energy resources and waste products. One question pertains to how fast microbiome perturbation due to URL construction wanes as microbiomes approach a pre-construction state over the following years. The Horonobe time series data indicate that the subsurface microbiomes are not undergoing rapid changes that could be indicative of rebound following laboratory construction in the five to nine years since the URL was constructed.

Any long term impact of the URLs on the subsurface should diminish with increasing distance from the access shafts and tunnels. Clustering of microbiomes based on organism presence/absence patterns generally is consistent with this prediction. The anomalously high abundance of SM1 archaea only close to the Horonobe tunnels may be explained by degassing of CO_2_ due to alteration of groundwater conditions. High CO_2_ conditions occur where this archaeon has been found elsewhere in high abundance (e.g., the CO_2_-driven Crystal Geyser, Probst et al. 2018).

The prevalence at Horonobe of *Methanoperedens*, an archaeon implicated in anaerobic methane oxidation, only at a site that is distant from the repository is interesting, given high levels of methane throughout the URL. Based on genomic information, methane oxidation is likely coupled to reduction of ferric iron (Hernsdorf et al., 2017; Nishimura et al., 2023). The sample was collected at the terminus of a downward inclined borehole where iron may be liberated from clay that accumulates there (Nishimura et al., 2023). Supporting the possibility that clay could stimulate growth of *Methanoperedens*, only the groundwater sampled from this site contained muddy suspended solids.

An intriguing finding regarding the Mizunami microbiomes is the high representation of CPR bacteria and DPANN archaea. Consistent with many prior studies (e.g., Wrighton et al. 2012), their gene inventories indicate that they are unlikely to be capable of living independently. CPR bacteria and DPANN archaea may be abundant in Mizunami high groundwater flow zones because low porosity enables selective mobilization of only small cells from more complex consortia attached to rock surfaces. A similar conclusion was suggested previously to explain the very high representation of CPR bacteria and DPANN archaea in the Crystal Geyser system (Probst et al., 2018).

We noted many differences in the taxa associated with the granitic vs. sediment-associated microbiomes that likely relate to the rock mineralogy. Granitic minerals such as biotite (iron, manganese), amphibole (iron, manganese), ilmenite (iron), chlorite (iron) (Yuguchi et al., 2010; 2018; Iwatsuki et al., 1999; Ishibashi & Yuguchi., 2017), Fe-, Mn-bearing calcite (Iwatsuki et al., 2000) and pyrite (e.g., iron, sulfur and selenium; Iwatsuki et al., 2002) may support microbial energy generation. Abiotic reactions involving minerals can be a source of methane and H_2_ (Stevens and McKinley, 1995), but carbon isotopic analyses of CH_4_ in the groundwater support a mixture of abiotic (Ino et al., 2017) and biogenic origin (Mills et al., 2010) and microbial consortia include methanogens. In the sedimentary rocks surrounding the Horonobe URL, buried organic matter, ferric iron oxides, iron-bearing clays (e.g., smectite) and pyrite may provide energy resources. High levels of dissolved CO_2_, H_2_, NH_3_ and CH_4_ in the groundwater are probably byproducts of microbial metabolism (Miyakawa et al., 2017). H_2_, NH_3_, and CH_4_ may persist in the Horonobe URL due to low availability of electron acceptors needed for the reactions that would consume them.

Given the possibility of Horonobe microbiome disruption close to the tunnels (e.g., Purkamo et al. 2018), communities sampled further from the tunnels may be more representative of the microbiomes surrounding the URL. Our data indicate that these communities are dominated by Deltaproteobacteria, especially *Syntrophobacterales*, which are predicted to be autotrophic sulfate-reducers that may also produce H_2_, supporting methanogenesis by abundant *Methanospirillum*. Betaproteobacteria such as *Methylophilales* are also relatively abundant in these samples and are implicated in methanol oxidation, the source of which may be methane oxidation linked to nitrate reduction, as performed by Gammaproteobacteria such as *Methylomonas* (Kits et al., 2015). Methanol oxidation is probably coupled to nitrate reduction given the lack of O_2_ (analogous to the process performed by *Methylomirabilis oxyfera;* Wu et al., 2015). The *Rhodocyclales* that are also abundant in these consortia are implicated in both nitrate reduction and sulfur oxidation.

A recent study provided insights into possible relationships between the composition of subsurface microbiomes and host rock type in the terrestrial subsurface (Soares et al. 2023). Despite low sample sizes resulting in a lack of statistical support, distinct lithologies were suggested to host distinct microbiomes, which is consistent with the taxonomic differences between the Horonobe and Mizunami URLs. The authors noted the widespread occurrence of Burkholderiales, Gammaproteobacteria and Clostridia, also groups represented in the URL microbiomes. However, we also note the prevalence of Parcubacteria (CPR bacteria), Deltaproteobacteria, Chloroflexi, Nitrospirae, and Bacteroidetes and numerous other groups, often candidate phyla (e.g., Altiarchiales and *Methanoperedens*). Thus, this study expands understanding of subsurface microbiome diversity, in part by providing information about species diversity and spatial variability.

Using the findings of the current study we can predict some impacts of the granite and sediment hosted microbiomes on geological disposal and storage. For example, proliferation of sulfate-reducing bacteria will generate sulfide that can corrode the metal containers potentially used to store radioactive waste (Pedersen, 2010; Stroes-Gascoyne, 2010; Enning and Garrelfs, 2014). It has been noted that production of sulfide by microbial communities in the terrestrial deep subsurface may be occurring even in the absence of geochemical evidence (Bell et al. 2020), highlighting the importance of microbial characterisation of these systems. Methanogens could also be problematic in that they can colonize the surfaces of steel containers and use the iron as the electron donor for methane production (Dinh et al., 2004; Mori et al., 2010; Hirano et al., 2022). Both sulfate reduction and methane production can be coupled to hydrogen metabolism, for which there are abundant genes. This raises the question of the extent to which hydrogen metabolizing microbes could impact subsurface H_2_ storage.

This study may be of interest beyond its immediate relevance to underground repositories or biodiversity and metabolism in granitic and sedimentary rock environments. Our analyses of microbial species overlap within subsurface locations (e.g., across depths of the Horonobe URL) and between subsurface locations on different continents may constrain rates of dispersal and genome mutation and calibrate intuition regarding global biodiversity (i.e., to what extent are the species or genera in every spatially separated underground site different?). The answer should depend on the degree of interconnection within the subsurface, the time since dispersal between distant locations, transit times from the surface to the subsurface, and evolutionary rates.

Considering first the case of within site dispersal, our results point to substantial genotypic overlap (i.e., very limited evolutionary divergence) within the Horonobe URL despite separation of the sampling sites by up to hundreds of meters of solid rock. This raises the question of whether genotype overlaps across sites can be explained by groundwater-mediated microbial transport. Comparison of d^18^O and dD between pore water and pumped groundwater indicates that there are “active” and “inactive” zones of groundwater flow within the Horonobe URL (Mochizuki et al., 2022). The groundwater samples collected for microbiology analyses are from “inactive” regions (**Fig. S6**). If measured hydraulic conductivities (**Table S2**) are predictive of transit times, microbial dispersal on thousands of years time scale might be possible. However, spatial heterogeneity in groundwater composition indicates extremely low interconnectivity, probably ruling out cell movement via water flow under the present hydraulic regime.

Alternatively, it is possible that microbial dispersal occurred under conditions that differ from the present. H and O stable isotopic measurements indicate that meteoric water was intruded into the Horonobe subsurface during glaciation (likely the Last Glacial Stage, between 12,000 and 70,000 years ago), when hydraulic gradients were higher than at present (Teramoto et al., 2010; Ishii, 2018; Nakata et al., 2018; Mochizuki & Ishii, 2022). Intrusion of meteoric water probably ceased when hydraulic gradients decreased due to sea level rise (Hanatani et al., 2010). If genomic overlap is due to microbial dispersal via groundwater flow >12,000 years ago, less than one mutation was fixed on average every ∼400 years in the Peregrinibacteria genome. We do not attribute genome consistency to inactivity because data indicate slow replication of these bacteria at the time of sampling. Some level of microbial activity is unsurprising, as an investigation of the deep terrestrial biosphere in the Äspö Hard Rock Laboratory in Sweden showed that subsurface populations are normally viable, with fast degradation of non-viable microbes likely resulting in the absence of populations that are not adapted to grow in oligotrophic, subsurface conditions (Lopez-Fernandez et al. 2018).

Assuming inter-site transport was possible, environmental heterogeneity could have precluded colonization by transported organisms. Thus, we attribute the genomic homogeneity within the Horonobe URL in part to consistent and stable geochemical and physical environments (as revealed by *in situ* measurements; Miyakawa et al., 2017; Sasamoto et al., 2015; Mochizuki & Ishii, 2022; 2023). The finding of biological stability is important for assessment of engineered subsurface environments, and thus long-term confinement of radioactive waste. Geological, hydrological, and geochemical stability for periods of at least tens of thousands of years is recommended for geological disposal (IAEA, 2003).

We detected closely related species overlaps between the Japanese URLs and underground locations in the USA. If inter-continental dispersal via air and/or water is relatively facile, and given the possibility of surface to Horonobe deep subsurface transport on the scale of tens of thousands of years, and in view of likely surface to subsurface groundwater movement rates at the USA locations (Supplementary Material), microbial dispersal may explain low levels of strain- to species level divergence in subsurface locations on two continents. These preliminary analyses were possible because large genome-resolved metagenomic datasets are available for several subsurface locations. Datasets of high quality genomes are being generated for many places around the world, and will enable further analyses of strain and species overlaps over geographically separated locations. Such research will advance our understanding of processes that structure the microbial biosphere, and by implication, the extent of microbial biodiversity in the Earth’s subsurface.

## Conclusions

This study was possible due to many years of planning and development to establish the two Japanese URLs as subsurface microbiology research sites. This is the first study to genomically evaluate the microbiomes in detail, over multiple years, and to compare the ecosystems in terms of community composition and microbiome function. The URL microbiomes are dominated by little known (i.e., candidate phyla) bacteria and archaea. Hydrogen, sulfur and nitrogen metabolisms are key to ecosystem survival in these dark, underground worlds. The research generated nearly comprehensive genomic datasets for organisms detected in both URLs, including for sites located near and far from the access tunnels, at multiple depths and over up to four years. The value of these genome collections for ecological, evolutionary, biotechnological and repository engineering studies will extend far beyond the current study. Our analyses provide answers to questions about microbiome stability and revealed surprising microbial community compositional and genotypic overlap over sites separated by hundreds of meters of rock, potentially explained by dispersal via slow groundwater flow or during a prior hydrological regime. Subsurface repositories for storage of H_2_ and radionuclides constructed in the future will benefit from insights regarding potential microbial impacts arising from microbial metabolism and constraints on the ways that the surrounding ecosystems will be impacted by repository engineering.

## Data availability

Prior to accession via NCBI, the draft genomes can be accessed via https://ggkbase.berkeley.edu/mizunami_genomes/organisms for the Mizunami URL and https://ggkbase.berkeley.edu/horonobe_genomes/organisms for the Horonobe URL. Some download functions may require you to sign up for a ggKbase account. Nine Horonobe datasets are publicly available via BioProject ID: PRJNA321556 (e.g., V-250m-2014 = Hor_250_2014). Upon publication, the sequences will be available via public data portals and the read datasets via the SRA.

## Acknowledgements

We thank Chris Greening for making available up to date hydrogenase HMMs and helpful discussion, Brian C. Thomas for bioinformatics support and helpful discussions, and Yusuke Watanabe, Mitsuru Kubota, Toshihiro Kato for groundwater sampling at the Mizunami Underground Research Laboratory. Funding for this project was provided by The Ministry of Economy, Trade and Industry of Japan, as “The project for validating near-field system assessment methodology in geological disposal” and “Development of the Technology for Integrating Radionuclide Migration Assessments” (2022 and 2023 FY, Grant Number: JPJ007597), the Japan Atomic Energy Agency (JAEA) Fund for Exploratory Researches (Houga fund), “Grant-in-Aid for Scientific Research (C) (Grant No. 19K05342) of the Japan Society for the Promotion of Science, National Science Foundation grant number OCE2049478 (to KA), A University of California, Berkeley Dissertation Year Fellowship (to LVA), The Berkeley Graduate Fellowship (to ALJ), Moore Foundation support (to JFB), NSF “Four Networks for Geologic Hydrogen Storage” grant 2230766 (to JFB), support from the Innovative Genomics Institute (to RS and JFB), the Watershed Function Scientific Focus Area funded by the U.S. Department of Energy, Office of Science, Office of Biological and Environmental Research under Award Number DE-AC02-05CH11231 (to AL) and sabbatical support from the University of California, Berkeley (to JFB).

## Author contributions

This study was designed by YA and JFB, with contributions from TI and HB. YA, HB, and YS obtained the metagenomic sequence datasets. RS and SL processed metagenomic data, RS, YA and JFB performed the binning. Organism representation and distribution patterns were analyzed by YA, RS and JFB. YA, JFB, ALJ, SD, CJC, and AL conducted the phylogenetic analyses. YA and KA took primary responsibility for metabolic analyses, TI, HB, KM, AM, EI, HM analyzed geochemical and hydrologic data. Hydrogenase analyses were performed by LVA, DG, YA and JFB. DG performed growth rate calculations. JFB performed genome curation. JFB and YA wrote the manuscript, with input from DG. All authors reviewed and commented on the manuscript prior to submission.

## Methods

### URL description

The Horonobe URL is located about 50 km south of Wakkanai in the northwestern peninsula of Hokkaido, Japan (Figure 1). It is situated in a low-lying coastal plain where Quaternary alluvium and terrace deposits overlie Tertiary and Cretaceous sediments that were deposited in the Mesozoic Tempoku Basin (Waseda et al., 1996). The Tempoku Basin is an on-shore basin that is elongated in the Horonobe area along a N-S axis. Seismic reflection surveys indicate that the current compressive E-W neotectonic stress to the west of the Horonobe area was established at around 2-3 Ma (Ogura and Kamon, 1992, Ito, 1999). Neogene strata in this area unconformably overlie Paleogene rocks. These strata consist of the Miocene Onishibetsu, Masuporo and Wakkanai formations, the Miocene-Pliocene Koetoi Formation and the Pliocene-Pleistocene Yuchi and Sarabetsu Formations (Iijima and Tada, 1981). All of these formations were deposited in a marine environment. The Koetoi and Wakkanai Formations are the main host rocks of the Horonobe URL. These formations are composed mainly of homogeneous siliceous rocks. The burial and subsidence of these formations occurred throughout the Neogene and Quaternary. Subsequent uplift and denudation started at about 1.3-1.0 Ma (Ishii et al., 2008). The Koetoi Formation is Neogene to Quaternary diatomaceous mudstones containing opal-A, and the Wakkanai Formation is Neogene siliceous mudstones containing opal-CT, with trace amount of quartz, feldspar, clay minerals, pyrite, calcite and siderite (Hiraga and Ishii, 2007; Ishii et al., 2007; Tachi et al., 2011). The Horonobe URL was constructed by JAEA to conduct basic geoscientific research and evaluate the feasibility and safety of geological disposal in deep sedimentary environments.

The Mizunami URL was located in the Gifu prefecture in central Japan (Figure 1). The URL was constructed by JAEA to conduct basic geoscientific research, but it was closed in 2019 as the research project was completed. Around the URL site, sedimentary rock (the Mizunami Group; 20-15 Ma, consisting of the Akeyo Formation) unconformably overlies Toki Granite (70 Ma). Toki granite is overlain by the Tertiary sedimentary rocks at ∼100 - 200 mbgl (meter below ground level) around the Mizunami URL. The Toki granite has three rock facies grading from muscovite-biotite granite, hornblende-biotite granite, and biotite granite. The constituent minerals are quartz, plagioclase, K-feldspar, biotite, hornblende, muscovite, accessory minerals and secondary minerals such as chlorite, calcite, and pyrite (Yuguchi et al., 2011, 2013). The Mizunami URL consisted of a main shaft, a ventilation shaft, sub-stages, and access tunnels at 300 and 500 m below ground level.

### Sample collection

Groundwater samples in the Mizunami URL were collected from six zones in four different boreholes (07MI07, 09MI20, 09MI21, 10MI26) at three depths 200, 300, 400 mbgl during 2014 and 2015 (Figure 1b; Table S1). Samples in the Horonobe URL were collected from five zones in three different boreholes (08E140C01, 07V140M03, 09V250M02) at five depths 140, 167, 194, 218, and 250 mbgl at the Horonobe URL during 2013 and 2016 (**Figure 1c**). All groundwater samples were obtained using multipacker systems (Nanjo et al, 2012). To minimize the influence of drilling and the installation of tools for hydrogeochemical monitoring, groundwaters were drained at least five times the section volume before monitoring for geochemical and microbial studies. The groundwater chemistry has been monitored since 2007, starting immediately after the drilling of these boreholes. The contamination of each groundwater sample by the drilling fluids was checked by measurement of the concentration of uranine and sodium naphthionate. The concentrations were below the detection limit. All but one sample for microbiological analyses was collected onto 0.22 μm pore size filters (type GVWP; Merck Millipore, Darmstadt, Germany) held in pressure-resistant stainless steel filter holders directly connected to the tubing outlet under in-situ hydraulic pressure conditions. The other sample (Hor_218_2014_10k) was collected using an ultrafiltration disc with 10,000 Da nominal molecular weight limit (type PLGC; Merck Millipore) after filtering through a 0.22 μm membrane filter. The volume of groundwater samples used for filtration was between 0.9 and 64 L at the Horonobe and 26 and 514 L at the Mizunami, depending on the cell densities in each groundwater sample (**Table S1**).

### DNA extraction and sequencing

DNA was extracted from the biomass collected on filters using the Extrap Soil DNA Kit Plus ver. 2 (Nippon Steel and Sumikin EcoTech Corporation, Tsukuba Japan). Genomic DNA libraries were prepared using TruSeq Nano DNA Sample Prep Kit (Illumina, San Diego, CA, USA) according to the manufacturer’s instructions. Quality of the library was examined using an Agilent 2100 bioanalyzer (Agilent Technologies) and paired-end 150-bp reads with a 550 bp insert size were sequenced by Hokkaido System Science Co., Ltd (Hokkaido, Japan), using an Illumina HiSeq 2500 (San Diego, CA, USA). DNA concentrations and sequencing information are presented in Table S1. Read datasets were assembled using IDBA_UD with the following parameters: -mink 40, -maxk 100, -step 20, and -pre_correction (Peng et al., 2012). Trimmed shotgun sequencing reads from each sample were mapped to all scaffolds >1000 bp, using Bowtie2 with default parameters (Langmead et al., 2009). For all scaffolds over 1000 bp, open reading frames were predicted with Prodigal using the meta setting (Hyatt et al., 2010). Functional annotations for all open reading frames were predicted using USEARCH (Edgar, 2010) searches against the Uniref100 (Suzek et al., 2007), Uniprot (Magrane and UniProt Consortium, 2011) and KEGG (Graham et al. 2018) to parse genes annotated with KEGG Orthology using Kofamscan (Aramaki et al. 2020), tRNA sequences were predicted using tRNAscan-SE (Schattner et al., 2005).

### Genome reconstruction

After assembly, scaffolds >1000 bp were binned by combination of phylogenetic profiles, read coverage and nucleotide content (GC proportion and tetranucleotide signatures) using ggKbase binning tools (https://ggkbase.berkeley.edu). Genomes that shared ≥ 95% average nucleotide identity were clustered to approximate species and the highest quality genome selected from each cluster. This analysis was performed using dRep (Olm et al. 2017).

### Organism distribution patterns

We conducted a census of organism types and abundances by identifying a non-redundant set of scaffolds encoding the ribosomal protein S3 (rpS3). This phylogenetically informative sequence, unlike 16S rRNA genes, tends to be well reconstructed from short read datasets (Olm et al. 2020). Reads were mapped to each scaffold encoding an rpS3 gene so that organism relative abundances in each sample could be determined. Species level groupings used rpS3 sequence comparisons with a ≥ 98% amino acid identity for classification of two organisms as approximately the same species.

### Phylogenetic analysis

The phylogeny of ribosomal protein S3 was calculated by aligning the sequences with select reference sequences using MUSCLE (Edgar, 2004). The alignments were manually trimmed in Geneious v.8 (BioMatters Ltd., San Francisco, CA, USA) to remove poorly aligned positions and columns composed of over 90% gaps before concatenation of protein sequences. Trees were built using RAxML v. 8.2.10 (Stamatakis, 2014) (as implemented on the CIPRES web server) (Miller et al., 2010), under the LG plus gamma model of evolution (PROTGAMMALG in the RAxML model section), and with the number of bootstraps automatically determined. RpS3 sequences were taxonomically classified using phylogenetic trees and NCBI protein BLAST searches. The in situ replication rates of microbial genomes were calculated using iRep v.1.1.14 (Brown et al. 2016) with default parameters.

### Functional gene analysis

Metabolic coding potential of all non-redundant sequences was explored by HMM searches against protein families downloaded from FunGene (Fish et al., 2013), TIGRFAM (Haft et al., 2003) and Pfam (Finn et al., 2014) as well as custom-built profile HMMs for several target genes (Eddy, 2011). Hydrogenases were annotated using profile hidden Markov model (HMM) searches (Eddy, 2011) with a custom set of [NiFe]-, [FeFe]- and [Fe]-hydrogenase HMMs (Søndergaard et al. 2016). Rubisco protein sequences were classified via phylogenetic analyses that used a custom dataset of reference sequences, including a subset from biochemically characterized types (**Data S2**).

### Similarity of organisms in each site

To evaluate the extent to which organisms present at each URL are now represented by genomes we identified all rpS3 genes from the combined dataset, regardless of whether or not they were in a draft genome bin and determined which samples contained sequences belonging to the dereplicated sequence clusters. Both dereplication and comparison of the dereplicated set to rpS3 sequences in each sample used an 99% nucleotide ANI threshold. This approach generated a presence/absence matrix that was sorted by the number of times each genome or unbinned sequence was detected across the sample series.

To evaluate how similar the collections of organisms represented by genomes from Mizunami and Horonobe were to each other and to groups of genomically defined organisms from other well sampled subsurface systems we first compared organisms from each site based on their rpS3 sequences encoded in their genomes, then based on average nucleotide identity (ANI) across alignable portions of their genomes. Different ANI cutoffs were used for different parts of the analysis. For the most closely related organisms, genome sequences and synteny were evaluated using Mauve (Darling et al., 2004). For one especially well assembled bacterial genome, manual curation was performed. This involved verification of overall assembly accuracy using mapped paired reads, removal of local assembly errors by excision of the portion not supported by reads and gap filling using reads and unplaced paired reads. Scaffold ends were extended using unplaced read pairs until circularization of the fully verified sequence was accomplished. The complete genome structure was evaluated using GC skew and cumulative GC skew using using gc_skew.py (https://github.com/christophertbrown/iRep) (Brown et al. 2016).

## Supplementary Information

### Mizunami and USA site hydrology constraints on cross-continent comparisons

The Rifle aquifer is shallow and impacted by inflow from the surface over weeks to months. Northern California groundwater wells respond to rainfall inputs over months. Although 10-20% of the Crystal Geyser water is derived from a Paleozoic aquifer, most is supplied by meteoric recharge (from a region 40-50 km away; Wilkinson et al., 2009). Residence time of approximately 9300 years for the groundwater was determined at the deeper part of sedimentary rocks overlying the Toki granite at the Mizunami URL (Iwatsuki et al., 2005), and the recent decrease in the water pressure following construction of the URL likely moved this water through the granitic URL (Hagiwara et al., 2015; Matsuoka & Hama, 2019). This groundwater intrusion provides a potential mechanism for microbial dispersal throughout the Mizunami URL. The case of Horonobe, results suggest that movement of surface water into the subsurface was possible ∼12,000 years ago. Thus, organisms could potentially have been distributed across all five locations ∼12,000 - 20,000 years ago.

### Supplementary Information

**Figure S1.**
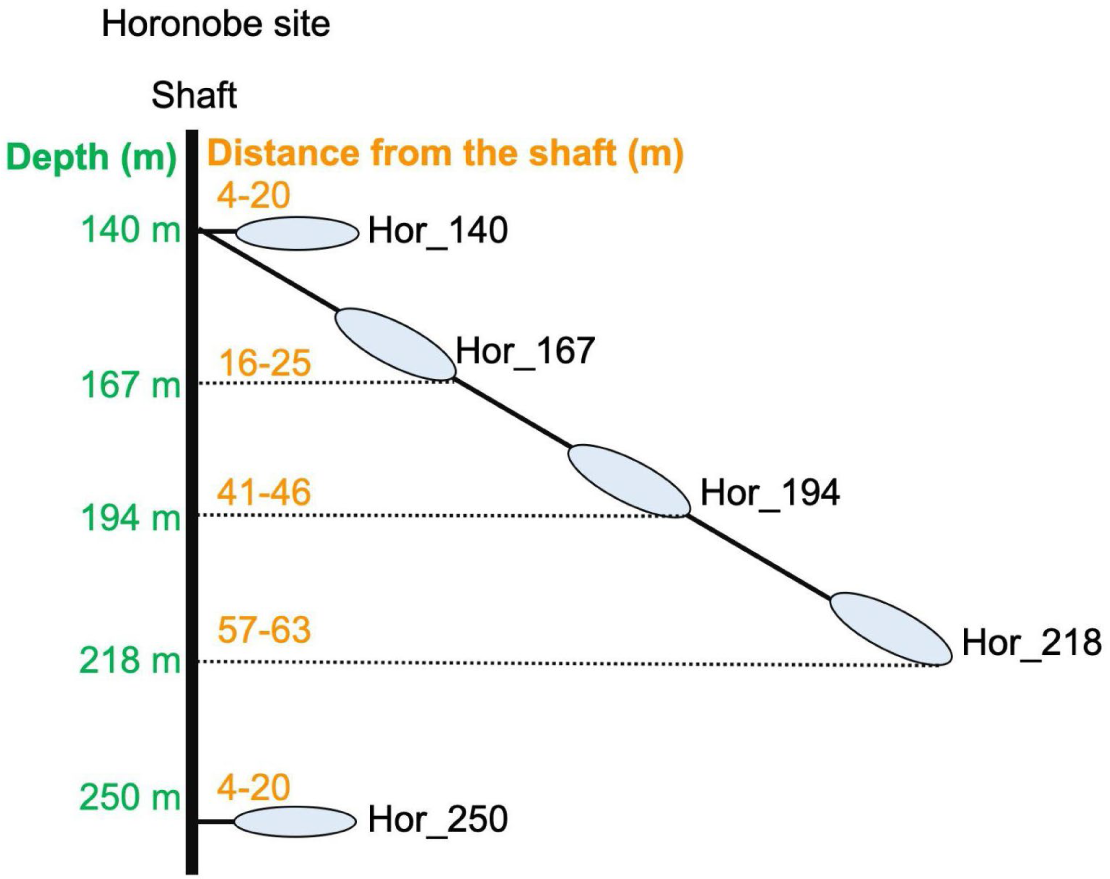
Diagram showing the layout of the Horonobe sampling sites. The y-axis indicates depth below the surface and the ovals indicate the distances that the groundwater sampling volumes are from the access tunnels.

**Figure S2.**
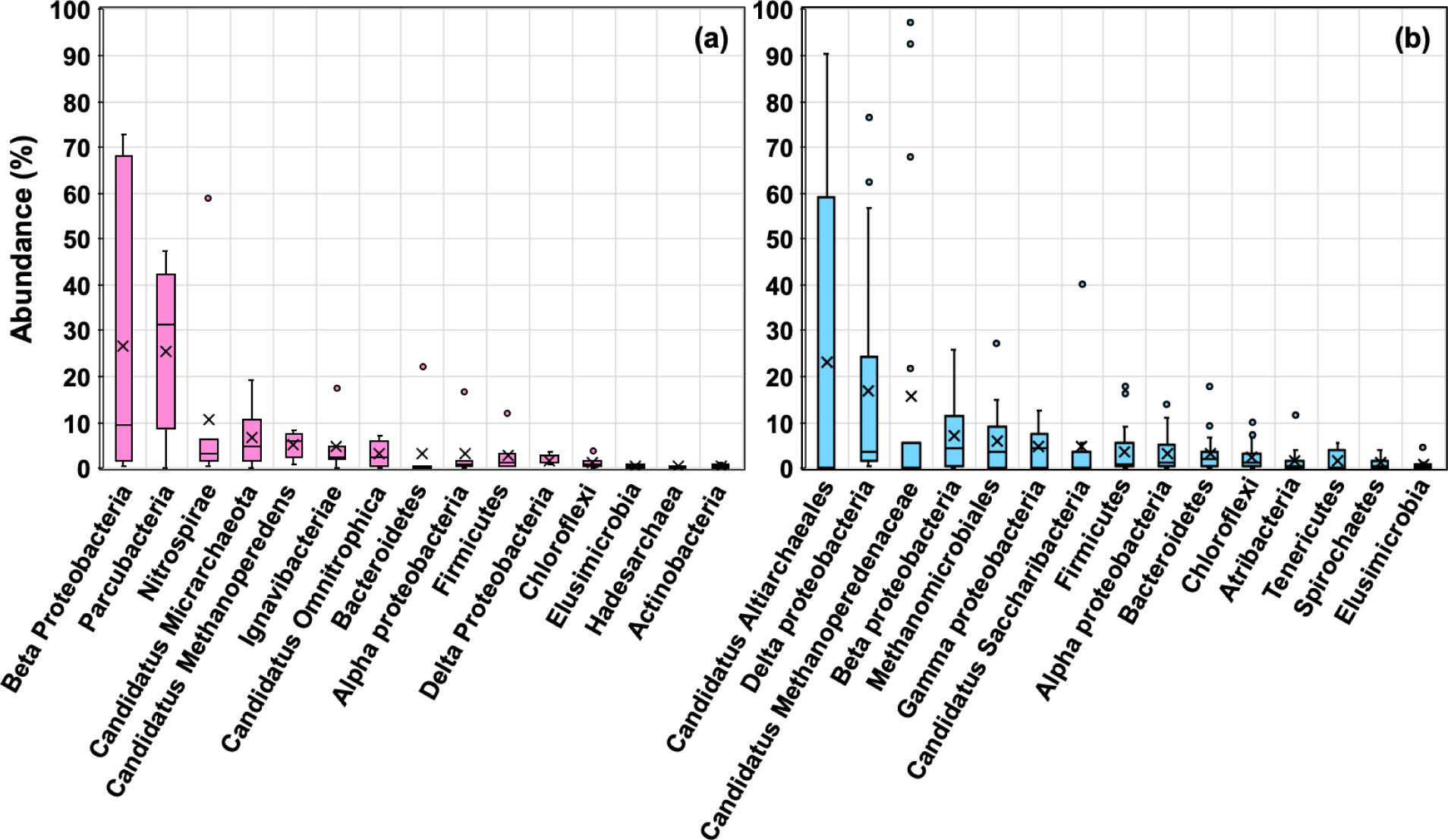
Relative abundances of the 15 most abundant organisms at the assigned rank based on normalized coverage values averaged over all samples. **(a)** Mizunami URL and **(b)** Horonobe URL. The edges of the box are the first and third quartile, crosses in the boxes are average proportions, circles are outliers.

**Figure S3.**
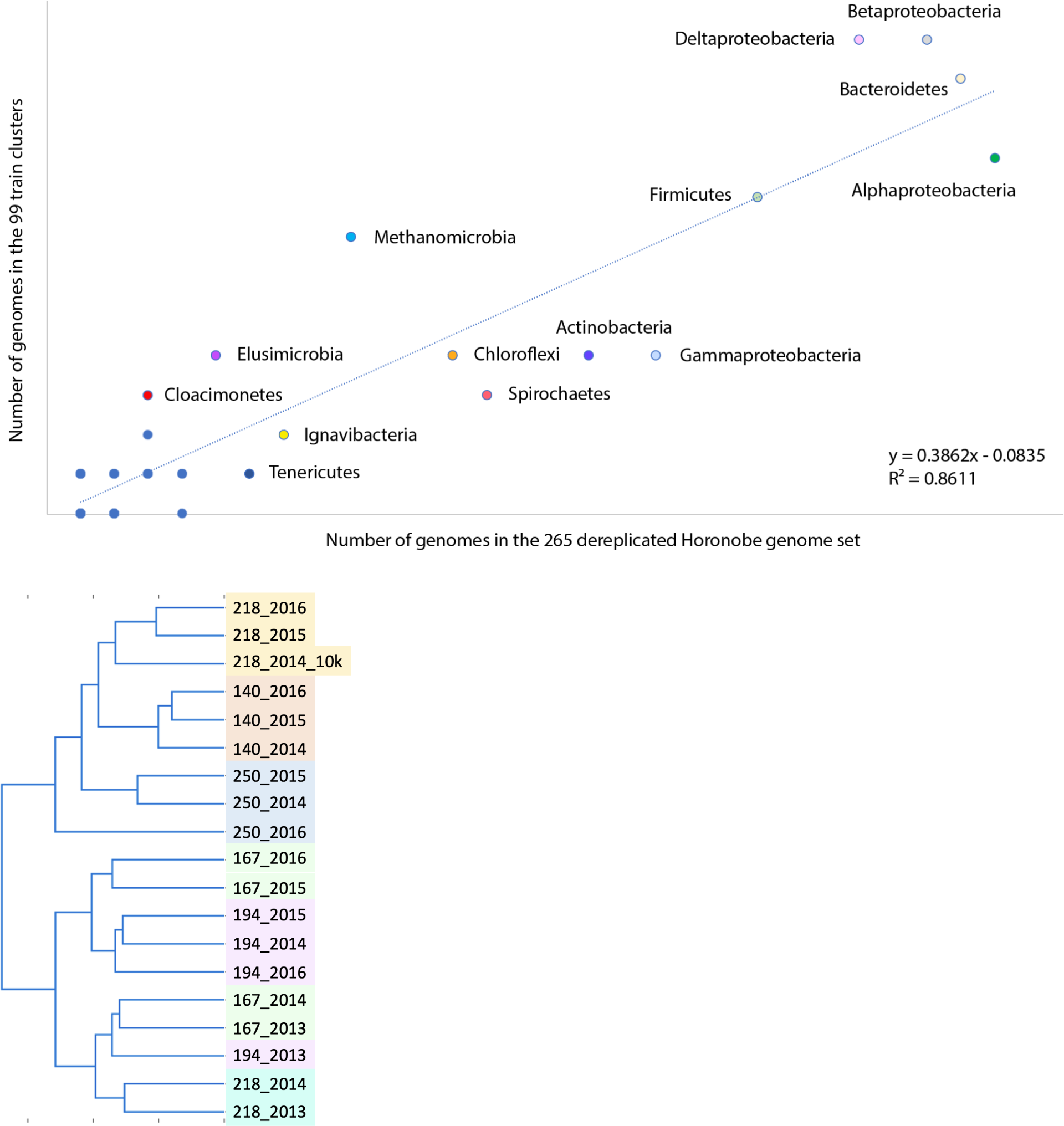
Analysis of the diversity structure and compositional similarity across sites and years for the Horonobe URL **(a)** The extent of within class-level diversity for each major taxonomic (approximately, phylum-level) group (dots) for 99 different species (40 groups represented by 265 genomes). Many points with low numbers of genomes assigned to a few classes are superimposed. Taxonomic affiliations are indicated for groups found at more than one site or depth (83% occur at > 1 depth). Generally, groups with many genomes (e.g., Alphaproteobacteria) have the genomes assigned to many classes (overall linear trend). **(b)** Hierarchical clustering based species presence/absence patterns for the Horonobe URL. Clustering mostly groups samples from the same site in different years. Samples from intermediate distances from the tunnels cluster with each other and with the earliest samples collected from the most distant site. Samples from 140 m and 250 m depths may be similar due to site locations very close to the access tunnels. The major subdivision in the sites is predicted by the presence of Candidatus groups Altiarchaeales, *Methanoperedens*, Saccharibacteria, *Deltaproteobacteria*, and *Betaproteobacteria*. Saccharibbacteria was detected in 2013 and 2014 at 167, 194 and 218 m depth sites, but not detected in 2015, contributing to the split in clustering for the 2015 samples.

**Figure S4.**
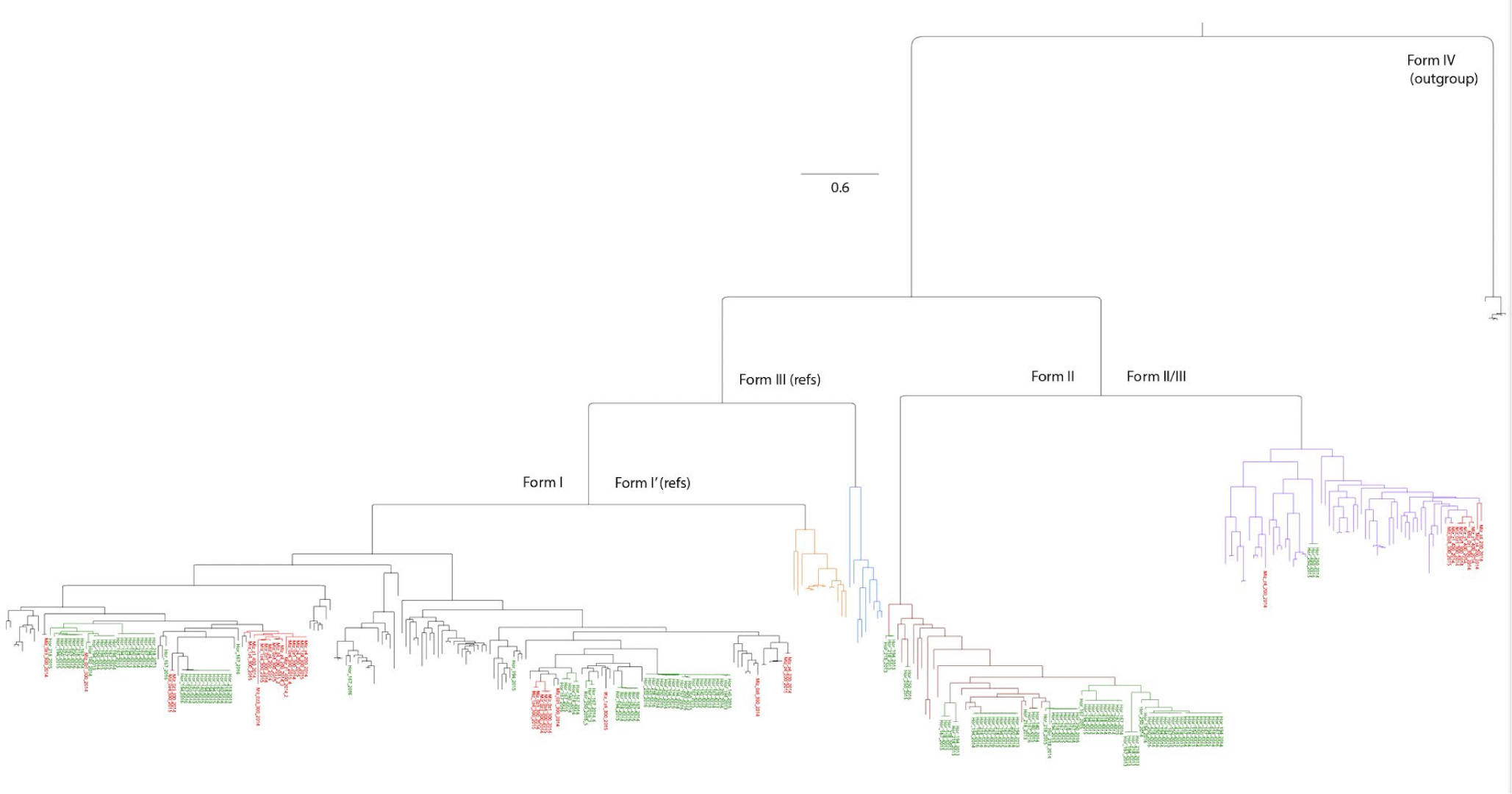
Phylogenetic tree showing the diversity of Rubisco forms I, II, and II/III. Forms I and II are likely involved in CO_2_ fixation via the CBB pathway. In most cases, phosphoribulokinase, another marker gene for this pathway, was identified (**Table S7**). Orange and blue branches are form 1’ and some form III sequences (included for context).

**Figure S5.**
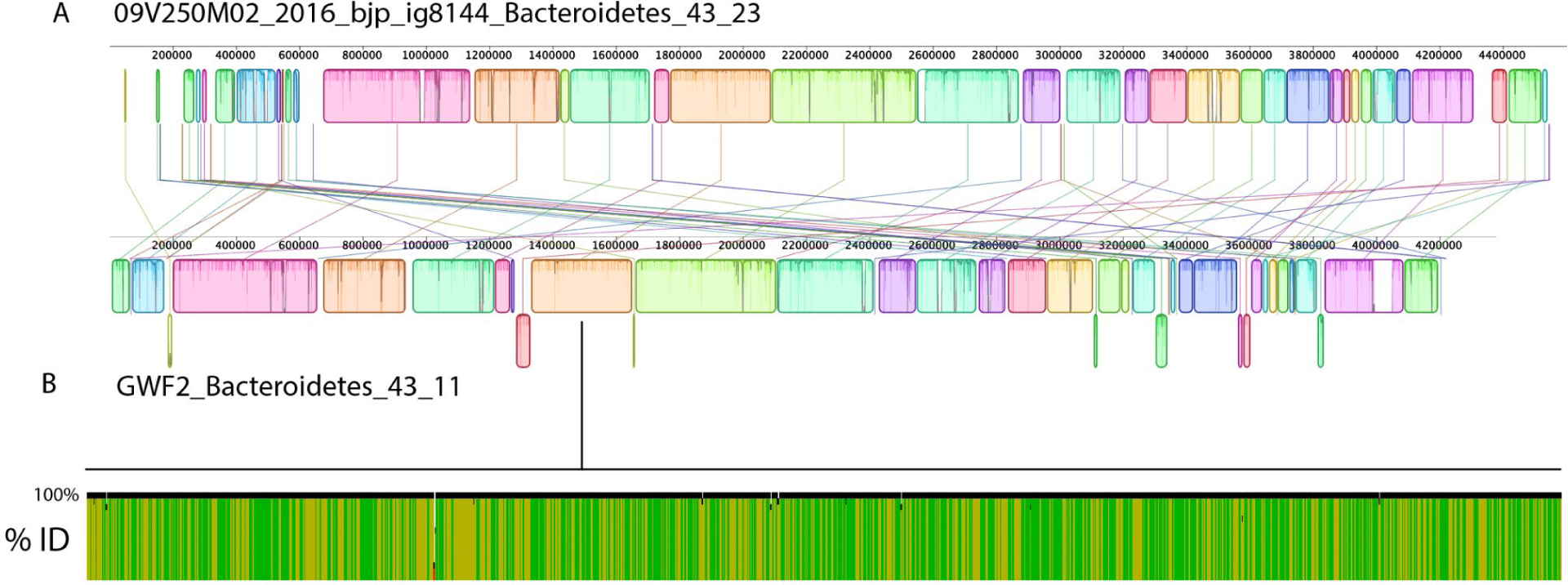
Comparison of the genome of a *Lentimicrobium* (Bacteroidetes) from the Horonobe URL (Hor_250_2016) to a genome from Rifle, CO groundwater (GWF2). Despite geographic and depth separation of these subsurface sites, the genome-wide average nucleotide identity determined based on alignable segments (which comprised > 85% of each genome bin) is 99.1%. (**a**) Alignment of the concatenated GWF2 *Lentimicrobium* genome bin scaffolds to the concatenated *Lentimicrobium* genome bin from Hor_250_2016. The colored blocks linked by lines indicate alignable sequence and the vertical axis of each box is % identity. (**b**) Segment of the aligned genomes showing nearly 100% sequence identity throughout.

**Figure S6.**
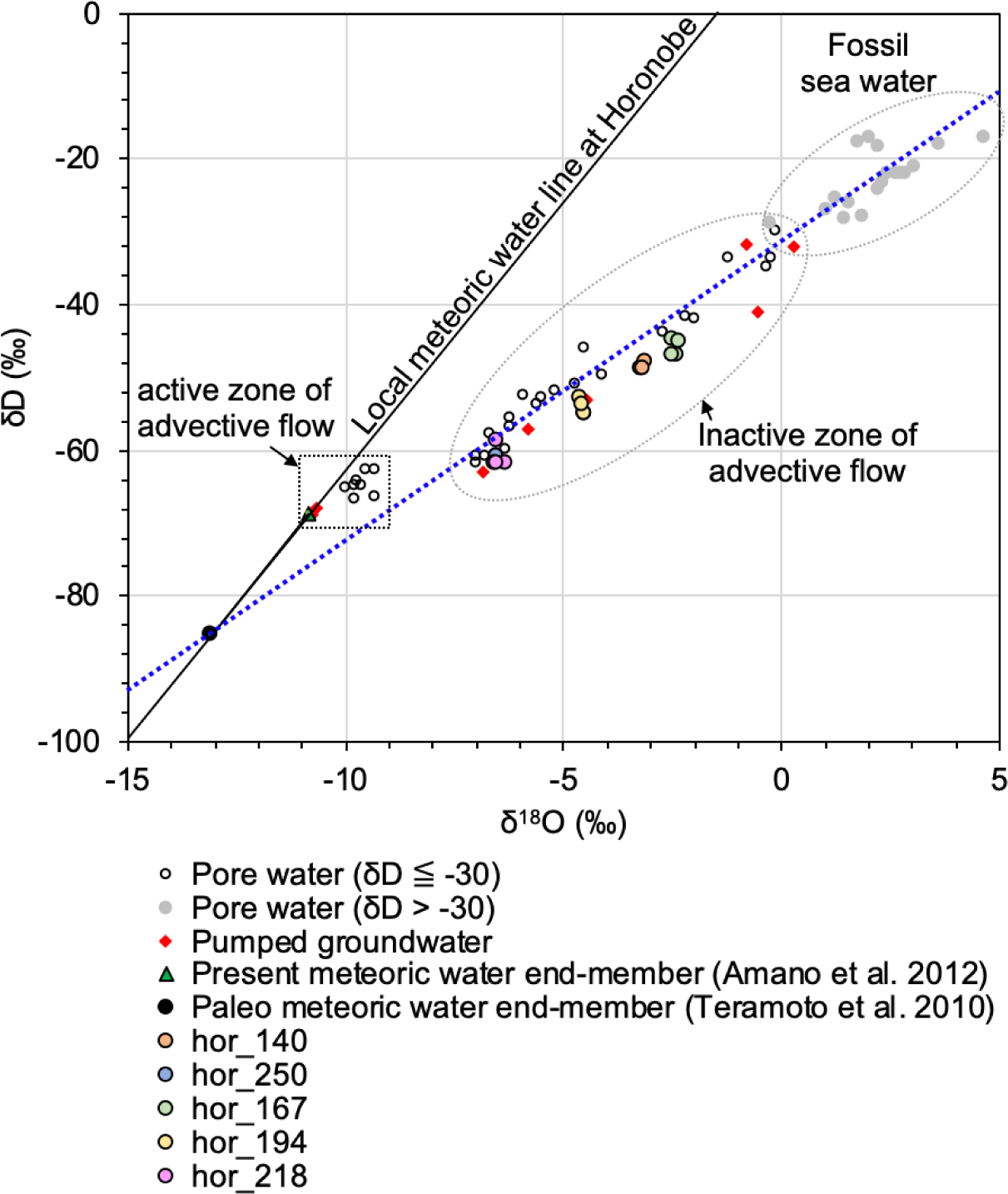
The relationship between d^18^O and dD in pore water and groundwater collected from the Horonobe URL (modified from Mochizuki & Ishii, 2022). Based on the comparison of d^18^O and dD between pore water and pumped groundwater, “active” and “inactive” zones of advective flow of meteoric water through fractures at present are suggested for the Horonobe samples (colored circles). The groundwater samples collected for microbiology analyses are classified as from currently inactive advective flow zones.

https://docs.google.com/spreadsheets/d/16RG_b07TiZx0E-U7qR5j7lZOsCo8WVQS/edit#gid=1381792296

### Supplementary Tables

https://docs.google.com/spreadsheets/d/16RG_b07TiZx0E-U7qR5j7lZOsCo8WVQS/edit#gid=1381792296

**Data S1. Non-redundant Newick format RPS3 protein phylogenetic tree**

(Data_S1_Newick format rpS3 protein phylogenetic tree.newick)

## References

Teske A, Biddle JF, Edgcomb VP, Schippers A. Deep subsurface microbiology: a guide to the research topic papers. Front Microbiol. 2013;4:1–3.

Hoshino T, Doi H, Uramoto GI, Wörmer L, Adhikari RR, Xiao N, Morono Y, D’Hondt S, Hinrichs KU, Inagaki F. Global diversity of microbial communities in marine sediment. Proc Natl Acad Sci USA. 2020;44:27587–97.

Pedersen K. Subterranean microorganisms and radioactive waste disposal in Sweden. Eng Geol. 1999;52:163–76.

Rajala P, Carpén L, Vepsäläinen M, Raulio M, Sohlberg E, Bomberg M. Microbially induced corrosion of carbon steel in deep groundwater environment. Front Microbiol. 2015;6:647.

Stroes-Gascoyne, S. & Hamon, C. & Maak, P. & Russell, S. The effects of the physical properties of highly compacted smectitic clay (bentonite) on the culturability of indigenous microorganisms. Applied Clay Science. 2010;47:155–162.

Stroes-Gascoyne S, West JM. Microbial studies in the Canadian nuclear fuel waste management program. FEMS Microbiol Rev. 1997;20:573–90.

Dopffel N, Mayers K, Kedir A, et al. Microbial hydrogen consumption leads to a significant pH increase under high-saline-conditions: implications for hydrogen storage in salt caverns. Sci Rep. 2023;13:10564.

Liu N, Kovscek AR, Fernø MA, Dopffel N. Pore-scale study of microbial hydrogen consumption and wettability alteration during underground hydrogen storage. Front Energy Res. 2023;11.

Zeng L, Sarmadivaleh M, Saeedi A, Chen Y, Zhong Z, Xie Q. Storage integrity during underground hydrogen storage in depleted gas reservoirs. Earth-Sci Rev. 2023;247:104625.

Wrighton KC, Thomas BC, Sharon I, Miller CS, Castelle CJ, VerBerkmoes NC, Wilkins MJ, Hettich RL, Lipton MS, Williams KH, Long PE, Banfield JF. Fermentation, hydrogen, and sulfur metabolism in multiple uncultivated bacterial phyla. Science. 2012;337:1661–65.

Anantharaman K, Brown CT, Hug LA, Sharon I, Castelle CJ, Probst AJ, et al. Metabolic handoffs shape biogeochemical cycles mediated by complex microbial communities. Nat Commun. 2016;7:13219.

He C, Keren R, Whittaker ML, et al. Genome-resolved metagenomics reveals site-specific diversity of episymbiotic CPR bacteria and DPANN archaea in groundwater ecosystems. Nat Microbiol. 2021;6:354–65.

Probst AJ, Ladd B, Jarett JK, Geller-McGrath DE, Sieber CMK, Emerson JB, Anantharaman K, Thomas BC, Malmstrom RR, Stieglmeier M, Klingl A, Woyke T, Ryan MC, Banfield JF. Differential depth distribution of microbial function and putative symbionts through sediment-hosted aquifers in the deep terrestrial subsurface. Nat Microbiol. 2018;3:328–36.

Hallbeck L, Pedersen K. Characterization of microbial processes in deep aquifers of the Fennoscandian Shield. Appl Geochem. 2008;23:1796–819.

Mehrshad M, Lopez-Fernandez M, Sundh J, Bell E, Simone D, Buck M, et al. Energy efficiency and biological interactions define the core microbiome of deep oligotrophic groundwater. Nat Commun. 2021;12:4253.

Konno U, Kouduka M, Komatsu DD, Ishii K, Fukuda A, Tsunogai U, et al. Novel microbial populations in deep granitic groundwater from Grimsel Test Site, Switzerland. Microbial Ecol. 2013;65:1–12.

Hernsdorf AW., Amano Y, Miyakawa K, Ise K, Suzuki Y, Anantharaman, K, Probst A., Burstein D., Thomas B.C., Banfield J.F. Potential for microbial H_2_ and metal transformations associated with novel bacteria and archaea in deep terrestrial subsurface sediments. The ISME Journal. 2017;11:1915–1929.

Ino K, Hernsdorf A, Konno U, Kouduka M, Yanagawa K, Kato S, et al. Ecological and genomic profiling of anaerobic methane-oxidizing archaea in a deep granitic environment. ISME J. 2017;12:31–47.

Iwatsuki T, Furue R, Mie H, Ioka S, Mizuno T. Hydrochemical baseline condition of groundwater at the Mizunami underground research laboratory (MIU). Applied Geochemistry. 2005;20:2283–2302.

Hayashida K, Munemoto T, Aosai D, Inui M, Iwatsuki T. Hydrochemical Investigation at the Mizunami Underground Research Laboratory -Compilation of Groundwater Chemistry Data in the Mizunami Group and the Toki Granite- (Fiscal Year 2014). JAEA-Data/Code 2016-001 (Technical Report by Japan Atomic Energy Agency). 2016. 10.11484/jaea-data-code-2016-001 (in Japanese with English abstract).

Amano Y, Yamamoto Y, Nanjyo I, Murakami H, Yokota H, Yamazaki M, et. Data of Groundwater from Boreholes, River Water and Precipitation for the Horonobe Underground Research Laboratory Project (2001-2010). JAEA-Data/Code 2011-023 (Technical Report by Japan Atomic Energy Agency). 2012. 10.11484/jaea-data-code-2011-023 (in Japanese with English abstract).

Sasamoto H, Yamamoto N, Miyakawa K, Mizuno. Data of Groundwater Chemistry Obtained in the Horonobe Underground Research Laboratory Project (2011–2013). JAEA-Data/Code 2014-033 (Technical Report by Japan Atomic Energy Agency). 2015. 10.11484/jaea-data-code-2014-033 (in Japanese with English abstract).

Miyakawa K, Ishii E, Hirota A, Komatsu D.D, Ikeya K, Tsunogai U. The role of low-temperature organic matter diagenesis in carbonate precipitation within a marine deposit. Appl. Geochem., 2017;76: 218–31.

Tamamura S, Miyakawa K, Aramaki N, Igarashi T, Kaneko K. A proposed method to estimate in situ dissolved gas concentrations in gas-saturated groundwater. Groundwater. 2018;56(1):118–30.

Adam PS, Panagiotis S, Borrel G, Brochier-Armanet C, Gribaldo S. The growing tree of Archaea: New perspectives on their diversity, evolution, and ecology. ISME J. 2017;11.

Brown CT, Hug LA, Thomas BC, Sharon I, Castelle CJ, Singh A, Wilkins MJ, Wrighton KC, Williams KH, Banfield JF. Unusual biology across a group comprising more than 15% of domain Bacteria. Nature. 2015;523:208–11.

Greening C, Biswas A, Carere CR, et al. Genomic and metagenomic surveys of hydrogenase distribution indicate H2 is a widely utilised energy source for microbial growth and survival. ISME J. 2016;10:761–77.

Ma K, Schicho R, Kelly R, Adams MW. Hydrogenase of the hyperthermophile Pyrococcus furiosus is an elemental sulfur reductase or sulfhydrogenase: evidence for a sulfur-reducing hydrogenase ancestor. Proc Natl Acad Sci USA. 1993;90:5341–5344.

Sasamoto H, Sato H, Arthur RC. Characterization of mineralogical controls on ammonium concentrations in deep groundwaters of the Horonobe area, Hokkaido. J Geochemical Exploration. 2018;188:318–325.

Brown CT, Olm MR, Thomas BC, Banfield JF. Measurement of bacterial replication rates in microbial communities. Nat Biotechnol. 2016;34(12):1256–63.

Nishimura H, Kouduka M, Fukuda A, Ishimura T, Amano Y, Beppu H, et al. Anaerobic methane-oxidizing activity in a deep underground borehole dominantly colonized by Ca. Methanoperedenaceae. Environ Microbiol Rep. 2023;15(3):197–205.

Yuguchi T, Tsuruta T, Nishiyama T. Zoning of rock facies and chemical composition in the Toki granitic body, Central Japan. GKK. 2010;39:50–70 (in Japanese with English abstract).

Yuguchi T, Sueoka S, Iwano H, Izumino Y, Ishibashi M, Danhara T, et al. Position-by-position cooling paths within the Toki granite, central Japan: Constraints and the relation with fracture population in a pluton. Journal of Asian Earth Sciences. 2018;169:47–66.

Iwatsuki T, Yoshida H. Groundwater chemistry and fracture mineralogy in the basement granitic rock in the Tono area, Gifu Prefecture, Japan -Groundwater composition, Eh evolution analysis by fracture filling minerals-. Geochem Jour. 1999;33:19–32.

Ishibashi M, Yuguchi T. Development of New Method for Evaluating the Mineral Distribution and Mode: Quantitative Image Analysis Using the Elemental Maps Obtained by the Scanning X-ray Analytical Microscope. Journal of the Japan Society of Engineering Geology. 2017;58: 80–93.

Iwatsuki T, Xu S, Itoh S, Abe M, Watanabe M. Estimation of relative groundwater age in the granite at the Tono research site, central Japan. Nuclear Instruments and Methods in Physics Research Section B: Beam Interactions with Materials and Atoms. 2000;172:524–529.

Iwatsuki T, Satake H, Metcalfe R, Yoshida H, Hama K. Isotopic and morphological features of fracture calcite from granitic rocks of the Tono area, Japan: a promising palaeohydrogeological tool. Appl Geochem. 2002;17(9):1241–57.

Stevens TO, McKinley JP. Lithoautotrophic microbial ecosystems in deep basalt aquifers. Science. 1995;270(5235):450–5.

Mills CT, Amano Y, Slater GF, Dias RF, Iwatsuki T, Mandernack KW. Microbial carbon cycling in oligotrophic regional aquifers near the Tono Uranium Mine, Japan as inferred from δ13C and Δ14C values of in situ phospholipid fatty acids and carbon sources. Geochimica et Cosmochimica Acta. 2010;74(13):3785–805.

Miyakawa K, Ishii E, Hirota A, Komatsu DD, Ikeya K, Tsunogai U. The role of low-temperature organic matter diagenesis in carbonate precipitation within a marine deposit. Appl. Geochem. 2017;76:218–231.

Purkamo L, Kietäväinen R, Miettinen H, Sohlberg E, Kukkonen I, Itävaara M, Bomberg M. Diversity and functionality of archaeal, bacterial and fungal communities in deep Archaean bedrock groundwater. FEMS microbiology ecology, 2018;94(8):fiy116.

Kits K, Klotz M, Stein L. Methane oxidation coupled to nitrate reduction under hypoxia by the Gammaproteobacterium *Methylomonas denitrificans*, sp. nov. Type Strain FJG1. Environmental Microbiology. 2015;17(9):3219–32.

Wu ML, Wessels JC, Pol A, Op den Camp HJ, Jetten MS, van Niftrik L. XoxF-type methanol dehydrogenase from the anaerobic methanotroph “Candidatus Methylomirabilis oxyfera”. Appl Environ Microbiol. 2015;81(4):1442–51.

Soares A, Edwards A, An D, Bagnoud A, Bradley J, Barnhart E, et al. A global perspective on bacterial diversity in the terrestrial deep subsurface. Microbiology. 2023;169.

Pedersen K. Analysis of copper corrosion in compacted bentonite clay as a function of clay density and growth conditions for sulfate-reducing bacteria. J Appl Microbiol. 2010;108:1094–1104.

Enning D, Garrelfs J. Corrosion of iron by sulfate-reducing bacteria: new views of an old problem. Appl Environ Microbiol. 2014;80(4):1226–36.

Bell E, Lamminmäki T, Alneberg J, Andersson AF, Qian C, Xiong W, Hettich RL, Frutschi M, Bernier-Latmani R. Active sulfur cycling in the terrestrial deep subsurface. ISME J. 2020 May;14(5):1260–72.

Dinh HT, Kuever J, Mußmann M, Hassel AW, Stratmann M, Widdel F. Iron corrosion by novel anaerobic microorganisms. Nature. 2004;427:829–32.

Mori K, Tsurumaru H, Harayama S. Iron corrosion activity of anaerobic hydrogen-consuming microorganisms isolated from oil facilities. J Biosci Bioeng. 2010;110:426–30.

Hirano, S.-i.; Ihara, S.; Wakai, S.; Dotsuta, Y.; Otani, K.; Kitagaki, T.; Ueno, F.; Okamoto, A. Novel *Methanobacterium* Strain Induces Severe Corrosion by Retrieving Electrons from Fe^0^ under a Freshwater Environment. Microorganisms 2022;10:270.

Mochizuki, Akihito & Ishii, Eiichi. Assessment of the level of activity of advective transport through fractures and faults in marine deposits by comparison between stable isotope compositions of fracture and pore waters. Hydrogeology Journal. 2022;30:813–827.

Teramoto M, Shimada J, Kunimaru T. Understanding groundwater flow regimes in low permeability rocks using stable isotope paleo records in porewaters. In: Taniguchi M, Holman IP, editors. Groundwater Response to Changing Climate. Florida: CRC Press; 2010. p. 79–86.

Ishii E. Assessment of Hydraulic Connectivity of Fractures in Mudstones by Single-Borehole Investigations. Water Resources Research. 2018;54.

Nakata K, Hasegawa T, Oyama T, Ishii E, Miyakawa K, Sasamoto H. An Evaluation of the Long-Term Stagnancy of Porewater in the Neogene Sedimentary Rocks in Northern Japan. Geofluids. 2018;1–21.

Mochizuki A, Ishii E. Assessment of the level of activity of advective transport through fractures and faults in marine deposits by comparison between stable isotope compositions of fracture and pore waters. Hydrogeology Journal. 2022;30:813–827.

Hanatani I, Munakata M, Kimura H, Sanga T. An analytic investigation of periglacial topography in Horonobe region Hokkaido. J. Nucl. Fuel Cycle Environ. 2010;17:55–70. 10.3327/jnuce.17.55 (in Japanese with English abstract).

Lopez-Fernandez M, Broman E, Turner S, Wu X, Bertilsson S, Dopson M. Investigation of viable taxa in the deep terrestrial biosphere suggests high rates of nutrient recycling. FEMS Microbiol Ecol. 2018;94(8):fiy121.

Miyakawa K, Ishii E, Hirota A, Komatsu DD, Ikeya K, Tsunogai U. The role of low-temperature organic matter diagenesis in carbonate precipitation within a marine deposit. Appl Geochem. 2017;76:218–31.

Mochizuki A, Ishii E. Paleohydrogeology of the Horonobe area, Northern Hokkaido, Japan: Groundwater flow conditions during glacial and postglacial periods estimated from chemical and isotopic data for fracture and pore water. Appl. Geochem. 2023;155:105737.

International Atomic Energy Agency (IAEA) Scientific and technical basis for geological disposal of radioactive wastes. Technical reports series, 2003;TRS no. 413.

Waseda A, Kajiwara Y, Nishita H, Iwano H. Oil-source rock correlation in the Tempoku basin of northern Hokkaido, Japan. Organic Geochemistry. 1996;24:351–362.

Ogura N, Kamon M. The subsurface structures and hydrocarbon potentials in the Tenpoku and haboro area, the northern Hokkaido, Japan. J. Jpn. Assoc. Petroleum Technol. 1992;57:32–44 (in Japanese with English abstract).

Iijima A, Tada R. Silica diagenesis of neogene diatomaceous and volcaniclastic sediments in northern Japan. Sedimentology. 1981;28:185e200.

Ishii E, Yasue K, Ohira H, Furusawa A, Hasegawa T, Nakagawa M. Inception of anticline growth near the Omagari Fault, northern Hokkaido, Japan. J Geol Soc Japan. 2008;114(6):286–99.

Hiraga N, Ishii E. Mineral and chemical composition of rock core and surface gas composition in Horonobe Underground Research Laboratory Project (Phase 1). JAEA-Data/Code 2007-022 (Technical Report by Japan Atomic Energy Agency). 2007.

Ishii E, Hama K, Kunimaru T, Sato H. Change in groundwater pH by infiltration of meteoric water into shallow part of marine deposits. J Geol Soc Japan. 2007;113:41–50.

Tachi Y, Yotsuji K, Seida Y, Yui M. Diffusion and sorption of Cs+, I- and HTO in samples of the argillaceous Wakkanai Formation from the Horonobe URL, Japan: A clay-based modeling approach. Geochim Cosmochim Acta. 2011;75:6742–59.

Yuguchi T, Amano K, Tsuruta T, Danhara T, Nishiyama T. Thermochronology and the three-dimensional cooling pattern of a granitic pluton: An example of the Toki granite, Central Japan. Contrib Miner Petrol. 2011;162:1063–77.

Yuguchi T, Tsuruta T, Hama K, Nishiyama T. The spatial variation of initial ^87^Sr/^86^Sr ratios in the Toki granite, Central Japan: Implications for the intrusion and cooling processes of a granitic pluton. J Miner Petrol Sci. 2013;108:1–12.

Nanjo I, Amano Y, Iwatsuki T, Kunimaru T, Murakami H, Hosoya S, Morikawa K. Development of a groundwater monitoring system at Horonobe Underground Research Center. JAEA Research 2011-048 (Technical Report by Japan Atomic Energy Agency). 2012. 10.11484/jaea-research-2011-048 (in Japanese with English abstract).

Peng Y, Leung HCM, Yiu SM, Chin FYL. IDBA-UD: a de novo assembler for single-cell and metagenomic sequencing data with highly uneven depth. Bioinformatics. 2012;28:1420–28.

Langmead B, Trapnell C, Pop M, Salzberg SL. Ultrafast and memory-efficient alignment of short DNA sequences to the human genome. Genome Biol 2009;10:R25.

Hyatt D, Chen GL, LoCascio PF, Land ML, Larimer FW, Hauser LJ. Prodigal: prokaryotic gene recognition and translation initiation site identification. BMC Bioinformatics. 2010;11:1–11.

Edgar RC. Search and clustering orders of magnitude faster than BLAST. Bioinformatics. 2010;26(19):2460–61.

Suzek BE, Huang H, McGarvey P, Mazumder R, Wu CH. UniRef: comprehensive and non-redundant Uni-Prot reference clusters. Bioinformatics. 2007;23:1282–1288.

Magrane M, UniProt Consortium. UniProt Knowledgebase: a hub of integrated protein data. Database. 2011:bar009.

Graham ED, Heidelberg JF, Tully BJ. Potential for primary productivity in a globally-distributed bacterial phototroph. ISME J. 2018;12(7):1861–66.

Aramaki T, Blanc-Mathieu R, Endo H, Ohkubo K, Kanehisa M, Goto S, Ogata H. KofamKOALA: KEGG Ortholog assignment based on profile HMM and adaptive score threshold. Bioinformatics. 2020;36(7):2251–52.

Schattner P, Brooks AN, Lowe TM. The tRNAscan-SE, snoscan and snoGPS web servers for the detection of tRNAs and snoRNAs. Nucleic Acids Res. 2005;33:W686–W689.

Olm, M., Brown, C., Brooks, B. et al. dRep: a tool for fast and accurate genomic comparisons that enables improved genome recovery from metagenomes through de-replication. ISME J 11, 2864–2868 (2017). 10.1038/ismej.2017.126

Olm MR, Crits-Christoph A, Diamond S, Lavy A, Carnevali PBM, Banfield JF. Consistent metagenome-derived metrics verify and delineate bacterial species boundaries. mSystems. 2020;14(5):e00731–19. DOI: 10.1128/mSystems.00731-19.

Edgar RC. MUSCLE: multiple sequence alignment with high accuracy and high throughput. Nucleic Acids Res. 2004;32:1792–97.

Stamatakis A. RAxML version 8: a tool for phylogenetic analysis and post-analysis of large phylogenies. Bioinformatics. 2014;30:1312–13.

Miller MA, Pfeiffer W, Schwartz T. Creating the CIPRES Science Gateway for inference of large phylogenetic trees. In 2010 Gateway Computing Environments Workshop (GCE). 2010;1–8.

Fish JA, Chai B, Wang Q, Sun Y, Brown CT, Tiedje JM et al. FunGene: the functional gene pipeline and repository. Front Microbiol. 2013;4:00291.

Haft DH, Selengut JD, White O. The TIGRFAMs data-base of protein families. Nucleic Acids Res. 2003;31:371–373.

Finn RD, Bateman A, Clements J, Coggill P, Eberhardt RY, Eddy SR et al. Pfam: the protein families database. Nucleic Acids Res. 2014:42:D222–D230.

Eddy SR. Accelerated profile HMM searches. PLoS Comput Biol. 2011;7(10):e1002195.

Søndergaard D, Pedersen CN, Greening C. HydDB: A web tool for hydrogenase classification and analysis. Sci Rep. 2016;6:34212.

Darling AC, Mau B, Blattner FR, Perna NT. Mauve: multiple alignment of conserved genomic sequence with rearrangements. Genome research. 2004 Jul 1;14(7):1394–403.

Olm, M.R., Brown, C.T., Brooks, B., and Banfield, J.F. dRep: A tool for fast and accurate genome de-replication that enables tracking of microbial genotypes and improved genome recovery from metagenomes. The ISME Journal. 2017;11:2864–68. doi: 10.1038/ismej.2017.126.

Wilkinson M, Gilfillan MV, Haszeldine RS, Ballentine CJ. Plumbing the depths: Testing natural tracers of subsurface CO_2_ origin and migration, Utah, USA. In: Grobe M, Pashin JC, Dodge RL. (Eds.), Carbon dioxide sequestration in geologic media -State of the science. American Association of Petroleum Geologists, AAPG Studies in Geology. 2009;59:619–634.

Hagiwara H, Iwatsuki T, Hasegawa T, Nakaga K, Tomioka Y. Shallow groundwater intrusion to deeper depths caused by construction and drainage of a large underground facility. Journal of Japanese Association of Hydrological Sciences. 2015;45:21–38

Matsuoka T, Hama K. Mizunami Underground Research Laboratory Project -Synthesis report on the R & D concerning important issues-. JAEA-Research 2019-2019-012. (Technical Report by Japan Atomic Energy Agency). 2019. 10.11484/jaea-research-2019-012 (in Japanese with English abstract).

